# Projected dynamics of breeding habitat suitability for a steppe-land bird warrant anticipatory conservation actions

**DOI:** 10.1101/2021.10.04.462847

**Authors:** Andrea Simoncini, Samuele Ramellini, Alexis Martineau, Alessandro Massolo, Dimitri Giunchi

**Author notes:** A. Simoncini and S. Ramellini should be considered as joint first authors. Corresponding author: Dimitri Giunchi, Dipartimento di Biologia, Università di Pisa, via Volta 6, 56126 Pisa (PI), Italy.

## Abstract

Understanding spatial and temporal variations of habitat suitability is fundamental for species’ conservation under global change. Steppic species are particularly sensitive to anthropogenic change and have undergone large declines in the last decades. We aimed to describe current and future breeding habitat suitability for the Eurasian stone-curlew *Burhinus oedicnemus*, a steppic species of conservation concern, and to identify critical areas for its conservation. We collected 1628 presence records covering the period 1992-2016. We developed a species distribution model using a dynamic Maxent algorithm and a set of pseudo-absences with a spatial density weighted on a fixed kernel density estimated on the presences, to mitigate the potential sampling bias. We projected this model under a set of carbon emission, socioeconomic and land-use/land-cover scenarios for the years 2030, 2050, 2070 and 2090. Finally, we described the cell-wise and mean change of breeding habitat suitability through consecutive time intervals and identified the areas critical for the species’ conservation.

All scenarios predicted a short-term northward shift of suitable areas, followed by a period of stability. We found no consistent trends in the mean change of breeding habitat suitability, and similar extents of suitable areas under current and future scenarios. Critical areas for the conservation of the species are mainly located in Northern Europe, Israel and parts of North Africa, the Iberian Peninsula and Italy. According to our results, the Eurasian stone-curlew has the potential to maintain viable populations in the Western Palearctic, but dispersal limitations might hinder the colonization of shifted suitable areas. Targeted conservation interventions in the critical areas are therefore recommended to secure the future of the species under global change.

## INTRODUCTION

Human-driven environmental change has relevant impacts on the distribution and life-history of organisms (Parmesan, 2006; Chen *et al*., 2011) and their likelihood of extinction (Román-Palacios & Wiens, 2020). Several studies reported distributional shifts in animal species according to climate change (Vanderwal *et al*., 2013), physiological constraints (Root *et al*., 2003) and in response to land-use and land-cover modifications (LULC; Sala *et al*., 2000). In birds, habitat suitability is affected by both climate and LULC change (Barbet-Massin, Thuiller, & Jiguet, 2011), and its variation has been linked to population trends (Green *et al*., 2008).

Steppe-land birds are particularly sensitive to environmental change, and large declines in steppe-land birds’ populations have been reported during the last decades, primarily as a consequence of agricultural intensification and afforestation following land abandonment (Burfield, 2005; Onrubia & Andrés, 2005). The Eurasian stone-curlew *Burhinus oedicnemus* (Linnaeus, 1758; hereafter stone-curlew) is a wide-ranging steppe-land species occurring in pseudo-steppes and farmlands in the Palearctic (Vaughan & Vaughan-Jennings, 2005). The species suffered a severe population decline (>30%) in the second half of the 20th century, mainly due to agricultural intensification (BirdLife International, 2018). It is now classified as Least Concern by the IUCN (BirdLife International, 2021), but information on the status of its populations is limited and positive trends might be due to the increased monitoring effort (Gaget *et al*., 2019). Indeed, the species is still considered of European conservation concern (SPEC3; BirdLife International 2017), and ongoing declines are reported for many regions of its range (BirdLife International, 2017; Gaget *et al*., 2019). Moreover, the stone-curlew is considered both an umbrella and flagship species (Caro, 2010; Hunter *et al*., 2016), and its conservation can therefore benefit steppe-land ecosystems (Hawkes *et al*., 2019).

To establish effective conservation practices, state-of-the-art habitat suitability scenarios for the focal species are required (Miller-Rushing, Primack, & Sekercioglu, 2010; Mcshea, 2014), and rigorous analyses of suitability dynamics are essential to reveal the sensitivity of species to global change and to guide conservation efforts (Stiels *et al*., 2021). Moreover, recent studies are increasingly evidencing the importance of distinguishing between the effects of long-term vs. short-term environmental change (Reside *et al*., 2010; Milanesi, Della Rocca, & Robinson, 2020). Previous studies using species distribution models (SDM, Guisan & Zimmermann 2000) to model habitat suitability for steppe-land birds reported northern range shifts and the persistence of large suitable areas according to global change scenarios (Estrada *et al*., 2016; Kiss *et al*., 2020). Huntley *et al*. (2007) used SDMs to forecast the future distribution of the stone-curlew under climate change scenarios, highlighting a northward shift of suitable areas.

Their analysis was limited to a coarse scale (ca. 50 km), Europe, one future period (end of the 21^st^ century) and a single climate scenario. An in-depth analysis integrating LULC and a range of climatic scenarios might validate their conclusions and provide time-varying predictions for conservation practitioners.

Therefore, in this study we aimed to: (i) test whether the stone-curlew’s distribution is driven by long-term or short-term environmental conditions, (ii) describe the current availability of suitable breeding habitats for stone-curlew, (iii) provide a range of future breeding habitat suitability scenarios considering both climate and LULC changes, (iv) test the hypothesis of a Northern shift for the stone-curlew, at higher spatial resolution than Huntley *et al*. (2007) and on multiple time-steps, and (v) identify the critical areas for its conservation. We predicted that (1) our models would highlight a northern shift of suitable breeding areas for the stone-curlew, (2) large areas of the Western Palearctic would be suitable for the species under future scenarios. As the stone-curlew is a species of warm temperate areas, it is unlikely that mean breeding habitat suitability will drop rapidly with global warming, as this is fastest at northern latitudes (IPCC, 2013). Therefore, we also predicted a stable or increasing mean breeding habitat suitability for the species under future conditions.

## MATERIALS AND METHODS

### Spatial scale

To reduce the spatial extent while retaining sufficient environmental variation, we limited our analysis to the Western Palearctic (*sensu* Snow & Perrins, 1998). We thus excluded *B. o. harterti* for poor data availability, and the two Macaronesian subspecies (*B. o. distincus*, *B. o. insularum*), as these show distinctive responses to the environment, being genetically different (Mori *et al*., 2017), and have a distribution restricted to small oceanic islands (Vaughan & Vaughan-Jennings, 2005).

As spatial resolution, we considered as appropriate the finest grain available based on future LULC scenarios (i.e. ~ 5.5×5.5 km grid cells; Chen *et al*., 2020) as this cell size approximates the breeding home-range of the species (Caccamo *et al*., 2011; Hawkes *et al*., 2021), thereby representing a biologically meaningful scale.

### Species occurrences

We used occurrences for the breeding season (May-July; Vaughan & Vaughan-Jennings, 2005) for the period 1992-2016 retrieving data from eBird, Global Biodiversity Information Facility (GBIF), ornitho.it, ornitho.cat and xeno-canto.org as in previous SDM studies (Avalos & Hernández, 2015; Coxen *et al*., 2017; Engelhardt, Neuschulz, & Hof, 2020; Ramellini *et al*., 2020). We also included survey data from the British Trust for Ornithology (in agreement with BTO’s data policy), data for Greece from the Ornithotopos database and the second European Breeding Bird Atlas (hereafter EBBA) and 2×2 km surveys in Greece (in formal agreement with the Hellenic Ornithological Society), data for the Deux-Sèvres department from the Nature79 database provided by Alexis Martineau. For Northern Italy, the dataset was integrated with nest points opportunistically collected by Dimitri Giunchi in the period 2012-2018.

We used data with a maximum positional uncertainty of 5.5 km to fit the resolution of predictors (Guisan, Thuiller, & Zimmermann, 2017), retaining the first occurrence in chronological order (i.e. the one reported in the first year available for the cell) when more than one point occurred in the same cell. We cut observations on the species’ range to exclude vagrant individuals in non-breeding areas, developing a range shapefile (see Fig. 1) based on the BirdLife species distribution dataset (http://datazone.birdlife.org/species/requestdis), the EBBA1 (Hagemeijer & Blair, 1997) and the EBBA2 (Keller *et al*., 2020). We then applied a spatial filtering of the data to reduce spatial autocorrelation (Kramer-Schadt *et al*., 2013), using the R package ‘spThin’ (v. 0.2.0; Aiello-Lammens *et al*., 2015) with a minimum distance of 5.5 km between points. The final dataset comprised 1628 presences (Fig. 1).

**Figure 1.**
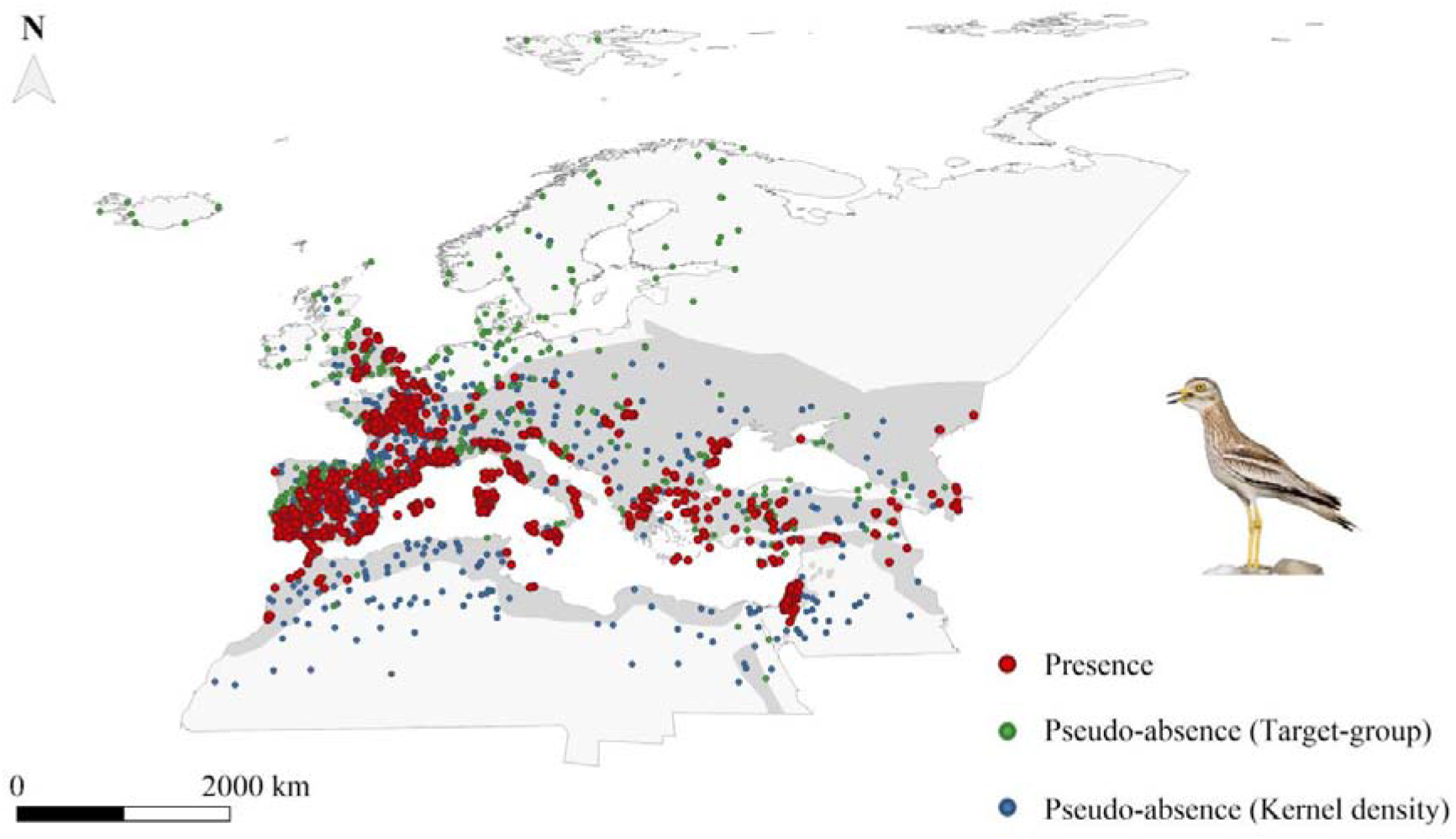
Occurrence and pseudo-absence points used to model breeding habitat suitability for the Eurasian stone-curlew. Target-group pseudo-absences are observations of other bird species recorded by the same observers of the presence data in the Global Biodiversity Information Facility (GBIF). Kernel density pseudo-absences are points with a spatial density weighted according to a fixed kernel density estimated from the presence points. For target-group and kernel density pseudo-absences, 500 randomly sampled points from the datasets are shown to improve image clarity. The grey area represents the species’ range in the Western Palearctic, with the exclusion of the Macaronesian archipelago. Photo by Saverio Gatto.

### Species distribution models

Contrasting results on the performance of different SDM algorithms have been reported (e.g. Qiao *et al*., 2015; Norberg *et al*., 2019). Therefore, we carried out preliminary tests to compare single-algorithm approaches and ensemble forecasting (i.e. a set of SDM algorithms combined to account for inter-model variability; Araújo & New, 2007). We then compared standard SDMs (hereafter static SDMs; Milanesi *et al*., 2020a) to dynamic SDMs. The first relate temporally dynamic occurrences to static predictor variables (e.g. 30-year climate averages), yielding potentially biased estimations of species-environment relationships (Milanesi *et al*., 2020a), whereas the latter combine occurrences with time-specific raster values (Reside *et al*., 2010; Milanesi *et al*., 2020a). We therefore tested whether the stone-curlew’s distribution is driven by long-term or short-term environmental conditions comparing static and dynamic SDMs. In summary, we developed and compared four classes of models:

i. single static SDMs: seven algorithms were developed (see Table S1 for a list of the employed techniques), using static variables (see *Environmental variables*);
ii. ensemble SDMs: the single static SDMs were combined according to the mean and the weighted average (Marmion *et al*., 2009). In the weighted average method, the AUCtest (see *Evaluation methods*) was used to weight model contribution to the final prediction. Static variables were employed;
iii. single dynamic SDMs: seven algorithms (Table S1) were employed, using dynamic variables (see *Environmental variables*);
iv. ensemble dynamic SDMs: the single dynamic SDMs were combined according to the mean and the weighted average, using dynamic variables.

All models were developed in the ‘biomod2’ (v. 3.3-7) R package (Thuiller *et al*., 2016).

### Pseudo-absence selection

We generated a set of pseudo-absence points (i.e. surrogates of true absences; Barbet-Massin *et al*., 2012), also using them as background points for Maxent. Pseudo-absences are often drawn at random in the study area, whereas occurrences typically show a spatial bias. Therefore, models could discriminate between the environmental features of sampled and unsampled areas, rather than between those of suitable and unsuitable areas (Phillips *et al*., 2009). To avoid this problem, Phillips *et al*. (2009) proposed the ‘target-group’ method, where pseudo-absences consist of other species’ data collected with the same methods used for the focal species. We used an approach similar to the observer-oriented one (Milanesi, Mori, & Menchetti, 2020). We considered as our pseudo-absences GBIF data of other bird species recorded by the same observers of the presence data, assuming they were collected with the same sampling bias. However, this approach could only be implemented on the GBIF data, and some biases might still arise. Therefore, we developed an alternative approach and refer to it as the ‘Kernel density approach’. We generated a large number of pseudo-absences, weighting their spatial distribution according to a fixed kernel density estimated on all the presence points, by means of the ‘adehabitatHR’ (v. 0.4.18) R package (Calenge, 2006). We used the *ad hoc* method for the estimation of the smoothing parameter (Worton, 1989). The procedure was independently repeated on the occurrences separated by year for the period 1992-2016. For both pseudo-absence selection methods, if a cell contained more than one record, the chronologically oldest was retained in the dataset. Finally, we removed pseudo-absences outside the study area and randomly sampled them to obtain a 0.1 prevalence (i.e. the ratio between presences and pseudo-absences ensuring an optimal performance; Barbet-Massin *et al*., 2012). To identify the best pseudo-absence method, we developed and evaluated a set of Maxent models for each method, using static variables (see *Environmental variables*). We expected the kernel density approach to outperform the target-group method, as the former might be more effective at cancelling out the potential spatial bias in the presence dataset.

### Environmental variables

We followed a two-step variable selection procedure, involving first an expert-based pre-selection and then a reduction of multicollinearity. In the first step, we used an expert-based approach (Santini *et al*., 2021) to derive 17 biologically meaningful environmental predictors (Table 1). We describe the biological rationale supporting expert-based variable choice and the process of variable preparation in the Appendix S1. For static SDMs, we averaged variables between the years 1973-2013 (climatic and ‘prey suitability’ variables) and 1992-2016 (LULC variables). Collectively, we refer to these variables as static variables. For dynamic SDMs, we associated occurrence/pseudo-absence pixels with year-specific values for a given variable (Milanesi *et al*., 2020a). We refer to these variables as dynamic variables. In the second step, we calculated the Variance Inflation Factor (VIF) to address multicollinearity between predictors. We used the *vifstep* function in the ‘usdm’ R package (v. 1.1-18; Naimi, 2013), using a threshold of three (Zuur, Ieno, & Elphick, 2010). We reported the initial and final VIF values for static and dynamic variables in Table 1. According to this analysis, the precipitation of the warmest quarter was only retained in the dynamic variable set and the temperature of the warmest quarter only in the static variable set. We excluded precipitation of the warmest quarter, retaining temperature of the warmest quarter for both static and dynamic variable sets. In the end, 13 variables were used in the models (Table 1).

**Table 1.** Candidate environmental variables to calibrate SDMs for the Eurasian stone-curlew. To describe the correlation among variables, we report the VIF (Variance Inflation Factor) for the initial and final sets of static and dynamic variables. Variables included in the dynamic Maxent model used for future projections are shown in bold, and their percentage contribution is reported.

| Variable | Static |  | Dynamic |  | Contribution (%) |
| --- | --- | --- | --- | --- | --- |
|  | VIF<sub>in</sub> | VIF<sub>fin</sub> | VIF<sub>in</sub> | VIF<sub>fin</sub> |  |
| **Percentage cover of agricultural areas (non-irrigated)** | 3.432 | 1.407 | 5.007 | 2.319 | 27.6 % |
| **Temperature of the warmest quarter** | 810.809 | 2.206 | 171.087 | - | 23.9 % |
| **Temperature seasonality** | 69.530 | 1.773 | 20.111 | 1.207 | 17.4 % |
| **Percentage cover of grass** | 1.736 | 1.274 | 2.541 | 1.684 | 10.8 % |
| **Percentage cover of agricultural areas (irrigated)** | 1.135 | 1.022 | 1.339 | 1.088 | 10.1 % |
| **Slope** | 1.463 | 1.229 | 1.977 | 1.684 | 9.0 % |
| **Percentage cover of trees** | 3.948 | 1.955 | 3.532 | 2.318 | 8.7 % |
| **Karst area** | 1.072 | 1.041 | 1.082 | 1.057 | 6.9 % |
| **Distance from water** | 2.002 | 1.831 | 1.767 | 1.557 | 4.8 % |
| **Percentage cover of urban areas** | 1.158 | 1.114 | 1.232 | 1.152 | 4.5 % |
| **Precipitation seasonality** | 1.747 | 1.607 | 1.823 | 1.722 | 3.6 % |
| **Percentage cover of shrubs** | 1.428 | 1.216 | 2.039 | 1.345 | 3.0 % |
| **Prey suitability** | 1.667 | 1.557 | 1.358 | 1.323 | 0.5 % |
| Annual temperature | 1000.396 | - | 142.475 | - | - |
| Annual precipitation | 6.717 | - | 3.686 | - | - |
| Precipitation of the warmest quarter | 7.323 | - | 3.754 | 2.198 | - |
| Percentage cover of bare areas | 9.983 | - | 6.365 | - | - |

#### Future variables

Future projections of bioclimatic conditions were obtained from four Intergovernmental Panel on Climate Change (IPCC) General Circulation Models (GCMs) that gained the minimum score in interdependence (Sanderson, Knutti, & Caldwell, 2015): ACCESS1-3, CESM1-BGC, CMCC-CM and MIROC5. We considered two emission scenarios corresponding to the IPCC’s Representative Concentration Pathways 4.5 and 8.5 (RCP 4.5 and RCP 8.5) for the years 2030, 2050, 2070 and 2090. Bioclimatic variables for every GCMxRCP combination and time interval were produced from the climatic variables available in the CHELSA CMIP-5 timeseries (Karger *et al*., 2017). Future LULC scenarios were retrieved from Chen *et al*., 2020. To create a LULC dataset consistent with bioclimatic scenarios, we selected LULC projections for the MIROC general circulation model (the remaining GCMs used for bioclimatic variables were not available), under RCP 4.5 and 8.5 and the Shared Socioeconomic Pathway 5, for 2030, 2050, 2070 and 2090. In future LULC scenarios, urban cover was kept at its 2016 value, due to the extremely low correspondence between the two LULC datasets.

### Model evaluation

We evaluated models based on the block cross-validation procedure implemented in the ‘ENMeval’ (v. 0.3.1) R package (Muscarella *et al*., 2014). In this procedure, data are partitioned into four geographically independent bins; four models are then produced, with three bins used for training and the remaining one used for testing. We used as performance statistics the area under the curve (AUC; Fielding & Bell, 1997) of the receiver operating characteristic plot, and the Continuous Boyce Index (CBI; Boyce *et al*., 2002). The AUC metric was computed with the R package ‘biomod2’ (v. 3.3-7, Thuiller *et al*., 2016); the CBI with the ‘ecospat’ (v. 3.1) R package (Di Cola *et al*., 2017). To detect overfitting in our models, we computed the difference between AUCtest and AUCtrain (AUCdiff; Radosavljevic & Anderson, 2014). In case of contrasting indications between AUCtest and CBI, we gave priority to the latter, as the usefulness of AUC in SDM has been questioned (Lobo, Jiménez-Valverde, & Real, 2008).

We evaluated variable importance using a model-independent approach (Thuiller *et al*., 2009) and expressed it as percentage. We then produced the response curves for each model using the evaluation strip method (Elith *et al*., 2005). To check the ecological validity of the models, we used an expert-based evaluation (Perennes *et al*., 2021). We defined two subjective scores: (i) Quality of Spatial Prediction (QSP) and (ii) Plausibility of Response Curve (PRC). The QSP was evaluated comparing qualitatively the spatial output of each model with the species’ distribution in the EBBA1 and EBBA 2 (Hagemeijer & Blair, 1997; Keller *et al*., 2020) and with the modelled distribution in the EBBA 2 (Keller *et al*., 2020). The PRC was evaluated based on the agreement between current knowledge on the species’ ecological preferences and the model’s response curves for the four variables with the highest percentage contribution. Both scores had three possible values (zero, one, two), with higher values indicating a better performance. Values were jointly discussed and assigned by Andrea Simoncini and Samuele Ramellini.

### Projections

The models developed during the model-selection phase were either projected on the static variables (static SDMs) or on variables representing the year 2016 (dynamic SDMs). The year 2016 was selected to describe current conditions as it was the closest year to the present time represented in our climatic dataset. The selected model was then projected to future conditions. We initially used a static LULC (i.e. the 2016 LULC variables) in future predictions to constrain climatic outputs (Pang, De Alban, & Webb, 2021). We then performed an additional analysis projecting models calibrated with the same historical LULC database on scenarios from the future dynamic LULC database (Chen *et al*., 2020). However, the difference between LULC databases did not allow quantitative interpretations of mean breeding suitability change.

In static LULC projections, for each RCP scenario and time interval we developed four projections (one per each GCM) averaging them in a final projection representing breeding habitat suitability for the species. We also computed the standard deviation between the four projections to account for the uncertainty arising from different climatic scenarios (Beaumont, Hughes, & Pitman, 2008; Porfirio *et al*., 2014). In dynamic LULC projections, we developed a single projection for each RCP and time interval, under the Shared Socioeconomic Pathway (SSP) 5 and the MIROC circulation model (other GCMs were not available).

When projected to new time periods, SDMs might encounter environmental conditions that are not found in the calibration area (Elith, Kearney, & Phillips, 2010; Zurell, Elith, & Schröder, 2012), producing spurious predictions (Owens *et al*., 2013). This occurs when: (i) a variable is outside the range found when training (i.e. strict extrapolation; Elith *et al*., 2010), or (ii) each variable is within the calibration range, but the combination of predictors is new (i.e. combinational extrapolation; Zurell *et al*., 2012). We thus used the environmental overlap mask (Zurell *et al*., 2012) to identify the areas where extrapolation occurs. We used the ‘mecofun’ (v. 0.1.0.9000) R package, with one bin per variable for strict extrapolation and five bins per variable to include combinational extrapolation (Zurell *et al*., 2012).

### Cell-wise change of habitat suitability

To describe the dynamics of breeding habitat suitability for the stone-curlew, we computed the difference of breeding suitability between consecutive time periods for each cell. Therefore, we obtained a set of rasters describing the magnitude of breeding habitat suitability change through consecutive time intervals, accounting for different RCPs and LULC scenarios.

### Change of mean habitat suitability

We evaluated the variation of breeding habitat suitability calculating the mean breeding habitat suitability (among all raster cells) for each future projection and considered its percentage change between two consecutive time periods, according to: each GCM under RCP 4.5 and RCP 8.5, and each RCP using the mean of all GCMs. This analysis was carried out for the static LULC projections only.

### Critical areas for conservation

Areas predicted as suitable under both current and future conditions are fundamental to plan *in situ* conservation actions (Estrada *et al*., 2016; Thuiller *et al*., 2019). We binarized each habitat suitability map according to the threshold that maximizes the sum of specificity and sensitivity (Liu, Newell, & White, 2016). Then, we summed all binarized projections and attributed a score of one if the cell was suitable in all the projections and a score of zero to all other cells. Additionally, we calculated the percentage of currently suitable cells representing critical areas.

Further details on the modelling procedure and workflow are reported in the Overview, Data, Model, Assessment and Prediction (ODMAP) protocol (Zurell *et al*., 2020; Appendix S1).

## RESULTS

### Pseudo-absence methods

The mean AUCtest, AUCdiff and CBI did not show marked differences between the two pseudo-absence methods (Table 2). However, except for the fourth run, the CBI was consistently higher for the kernel density approach, which also provided higher quality spatial predictions (Table 2, Fig. S1.2). Therefore, we decided to use the kernel density approach for subsequent analyses. We selected the first run due to the particularly high CBI, a fair AUCtest and a high-quality spatial prediction (Table 2).

**Table 2.** Evaluation metrics of the static Maxent models for the Eurasian stone-curlew calibrated with two pseudo-absence approaches (target-group and kernel density), for four cross-validation runs. We report Area Under the ROC (Receiver Operating Characteristic) Curve (AUC) of the training and test datasets and their difference; Continuous Boyce Index (CBI); Quality of Spatial Prediction (QSP) and Plausibility of Response Curve (PRC). Quality of Spatial Prediction (QSP) and Plausibility of Response Curve (PRC) are qualitative subjective scores (value of zero, one or two) describing the ecological validity of a model. Mean and standard error are reported for each approach.

| Target group |  |  |  |  |  |  |
| --- | --- | --- | --- | --- | --- | --- |
| Run | *AUC<sub>train</sub>* | *AUC<sub>test</sub>* | *AUC<sub>diff</sub>* | *CBI* | *QSP* | *PRC* |
| 1 | 0.891 | 0.679 | 0.212 | 0.852 | 1 | 2 |
| 2 | 0.878 | 0.746 | 0.132 | 0.977 | 0 | 2 |
| 3 | 0.854 | 0.824 | 0.030 | 0.842 | 1 | 2 |
| 4 | 0.850 | 0.832 | 0.018 | 0.654 | 1 | 2 |
| Summary |  |  |  |  |  |  |
| *Mean (SE)* | 0.868 (0.010) | 0.770 (0.036) | 0.098 (0.046) | 0.831 (0.067) | 0.75 (0.25) | 2 (0) |
| Kernel density |  |  |  |  |  |  |
| Run | *AUC<sub>train</sub>* | *AUC<sub>test</sub>* | *AUC<sub>diff</sub>* | *CBI* | *QSP* | *PRC* |
| 1 | 0.888 | 0.783 | 0.105 | 0.999 | 2 | 1 |
| 2 | 0.884 | 0.871 | 0.013 | 0.985 | 1 | 1 |
| 3 | 0.885 | 0.727 | 0.158 | 0.887 | 2 | 2 |
| 4 | 0.892 | 0.799 | 0.093 | 0.333 | 2 | 2 |
| Summary |  |  |  |  |  |  |
| *Mean (SE)* | 0.887 (0.002) | 0.795 (0.030) | 0.092 (0.030) | 0.801 (0.158) | 1.75 (0.25) | 1.5 (0.29) |

Dynamic models gained consistently higher CBI and generally higher QSP and PRC than static models (Table 3). Ensemble SDMs showed high AUCtest, but not significantly higher than the best-performing single methods (i.e. Random Forest, Maxent, GLM), and a discrete overfitting (Table 3). The CBI and PRC showed the same general pattern, with higher estimates for dynamic ensemble models (Table 3). Spatial predictions did not differ significantly between static and dynamic models, except for the ANN algorithm (Fig. S1.3-S1.4). However, static models identified suitable areas at the southern edge of the study area in the African continent where the species is absent (Keller *et al*., 2020). Instead, we found a higher agreement in these areas between dynamic models’ projections and the actual species’ range (Fig. S1.3-S1.4). Ensemble models gained good quality spatial outputs but tended to over-predict habitat suitability at high latitudes (e.g. in Arctic Russia, where the species is not found) and under-predict at lower latitudes (e.g. in France and the Iberian Peninsula, where the species is widespread). Instead, GAM and Maxent provided excellent projections in these areas (Fig. S1.3-S1.4). Therefore, we retained the best performing single dynamic model (i.e. Maxent; Fig. 2), that showed a good AUC (0.803), an excellent CBI (0.999) and maximum QSP and PRC values (Table 3).

**Table 3.**
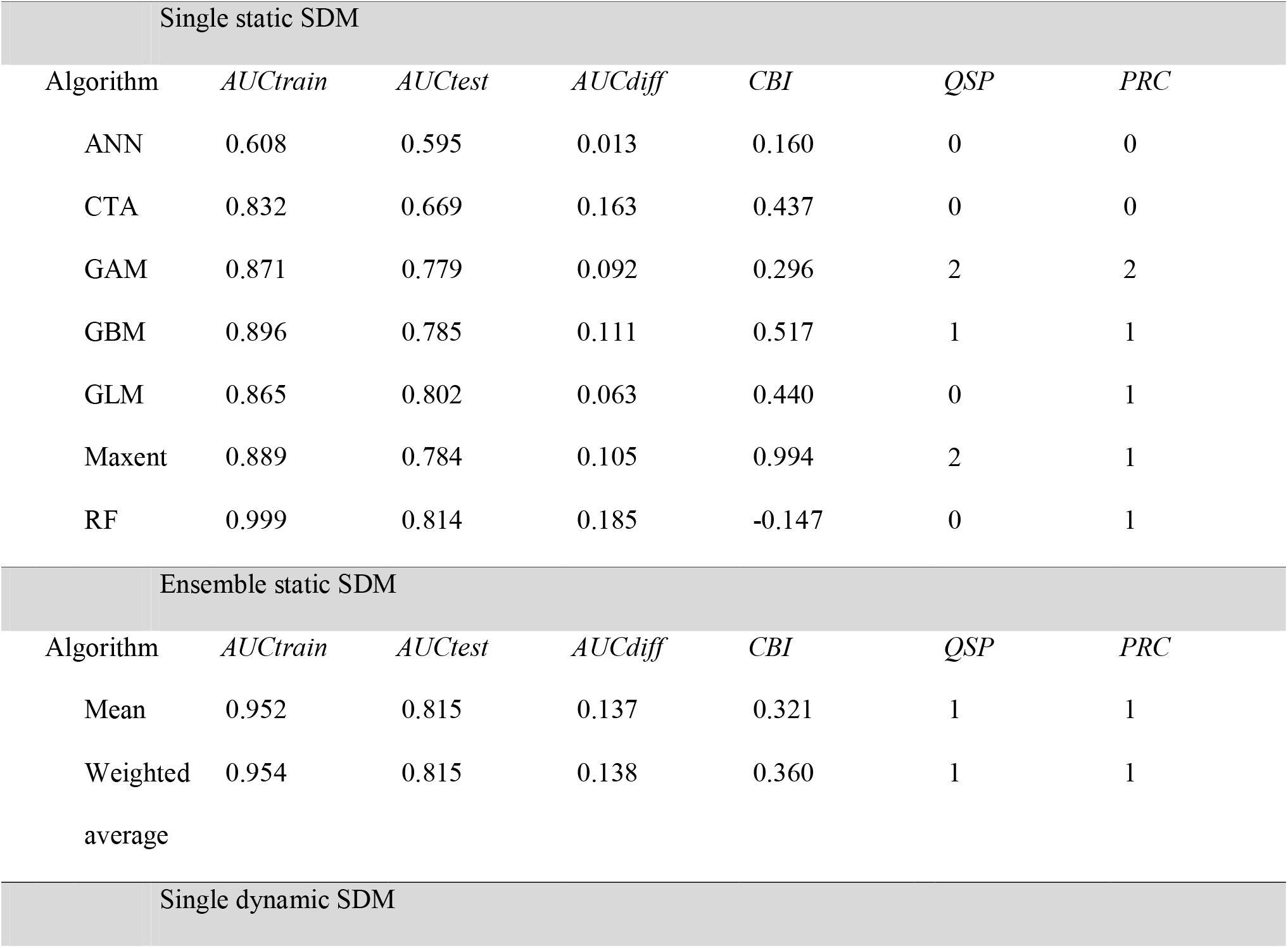
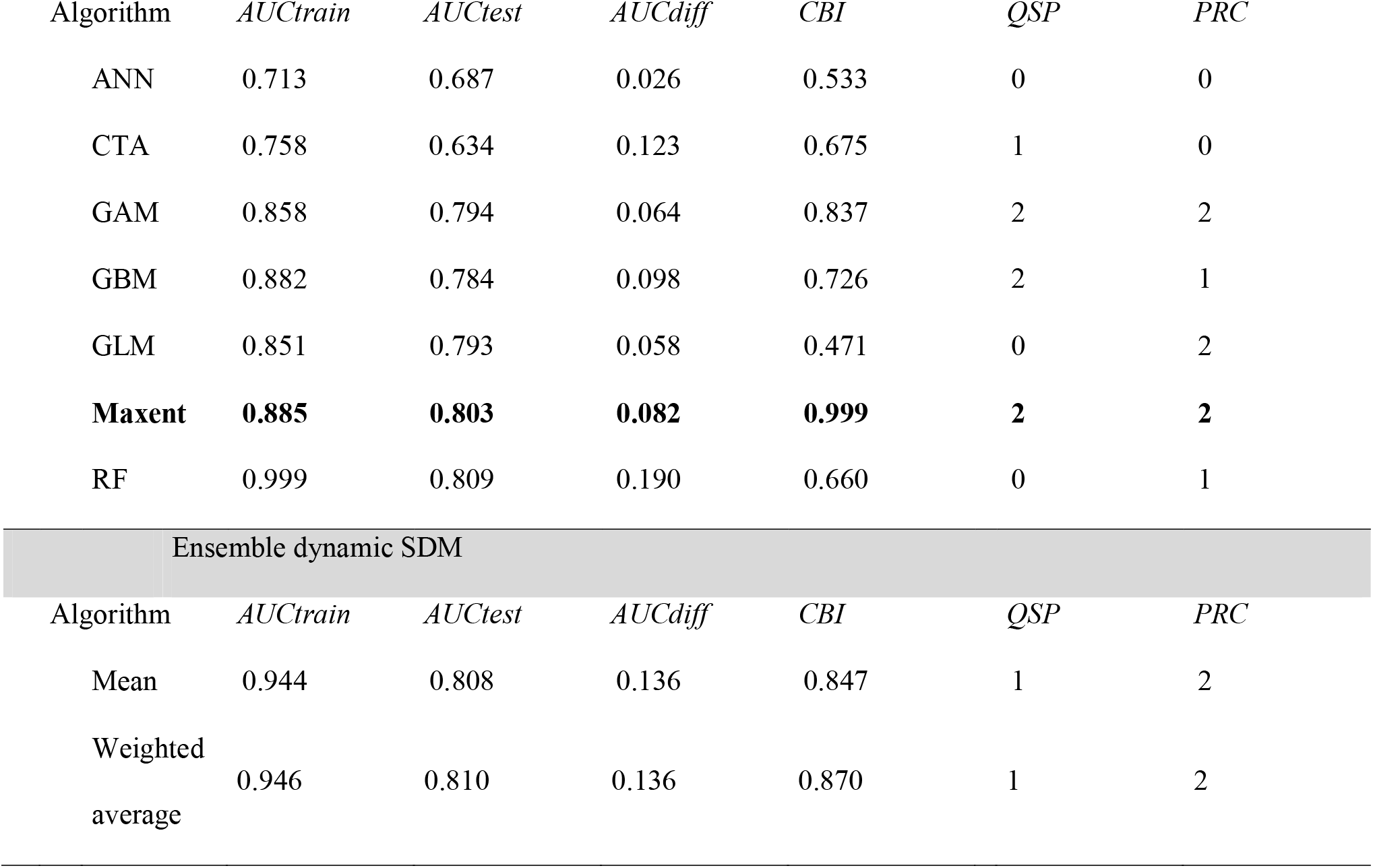
Evaluation metrics of four species distribution modelling strategies for the Eurasian stone-curlew: single static, ensemble static, single dynamic and ensemble dynamic. The following single algorithms have been used: Artificial Neural Networks (ANN), Classification Tree Analysis (CTA), Generalized Additive Models (GAM), Generalized Boosting Models (GBM), Generalized Linear Models (GLM), Maximum entropy (Maxent), Random Forest (RF). The following metrics are reported: Area Under the ROC (Receiver Operating Characteristic) Curve (AUC) of the training and test datasets and their difference; Continuous Boyce Index (CBI). Quality of Spatial Prediction (QSP) and Plausibility of Response Curve (PRC) are subjective scores (value of zero, one or two) describing the ecological validity of a model. The model used for future projections is highlighted in bold.

**Figure 2.**
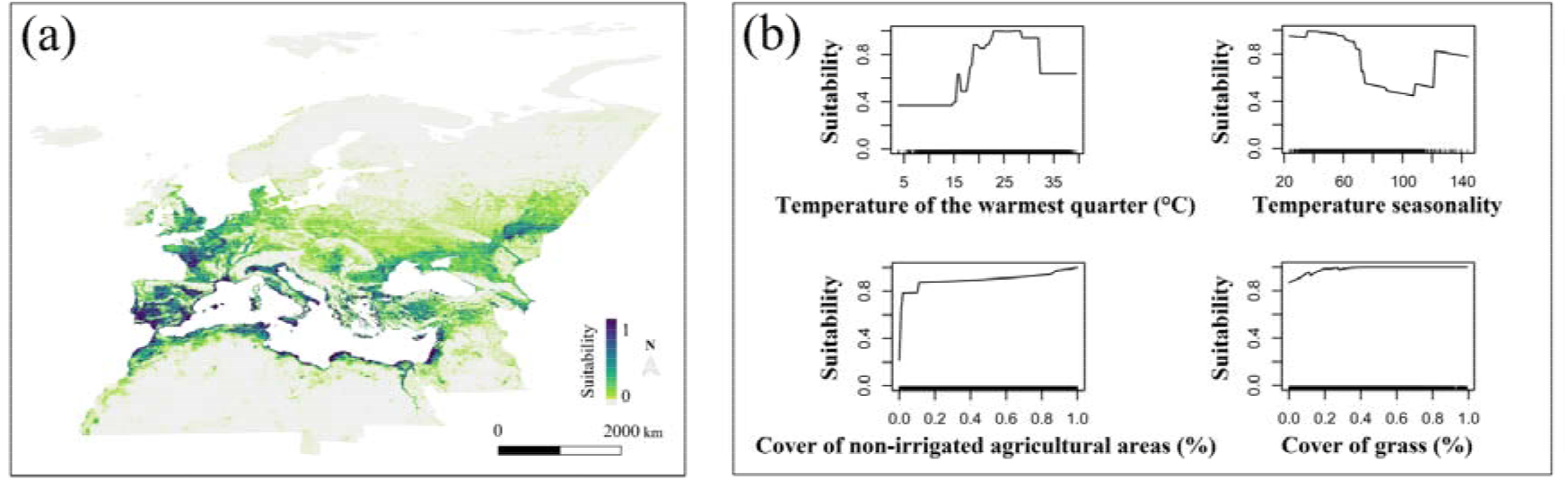
(a) Current breeding habitat suitability map according to the dynamic Maxent model selected to project breeding habitat suitability for the Eurasian stone-curlew under global change scenarios, and (b) response curves for the four environmental variables with the highest percentage contribution in the same model.

### Future projections

Projections of future breeding habitat suitability for the species differed between static and dynamic LULC scenarios, but not between RCPs and time periods (Fig. S1.5). Dynamic LULC projections described a similar spatial distribution of suitable areas for the species compared to static LULC projections, but with higher suitability estimates (Fig. S1.5). The agreement between GCMs for static LULC projections was mostly high (Fig. S1.7). Strict extrapolation was generally low (Fig. S1.8-S1.9, Fig. S1.12), although it occurred consistently in some regions at the margins of our study area (e.g. Northern Russia) and it was generally higher under RCP 8.5. Combinational extrapolation was far more extensive and showed increasing values with latitude and under RCP 8.5 (Fig. S1.10-S1.12).

### Cell-wise change of habitat suitability

The cell-wise change of habitat suitability was particularly evident in the first time interval (i.e. 2016-2030), while quite small in the subsequent ones (Fig. 3). Suitable areas for the species are expected to shift northwards. Broad areas in the central-southern part of the range are predicted to be lost, most of them in the time interval 2016-2030. Differences between static and dynamic LULC are marked, with the suitability of the latter predicted to undergo a higher change (Fig. 3).

**Figure 3.**
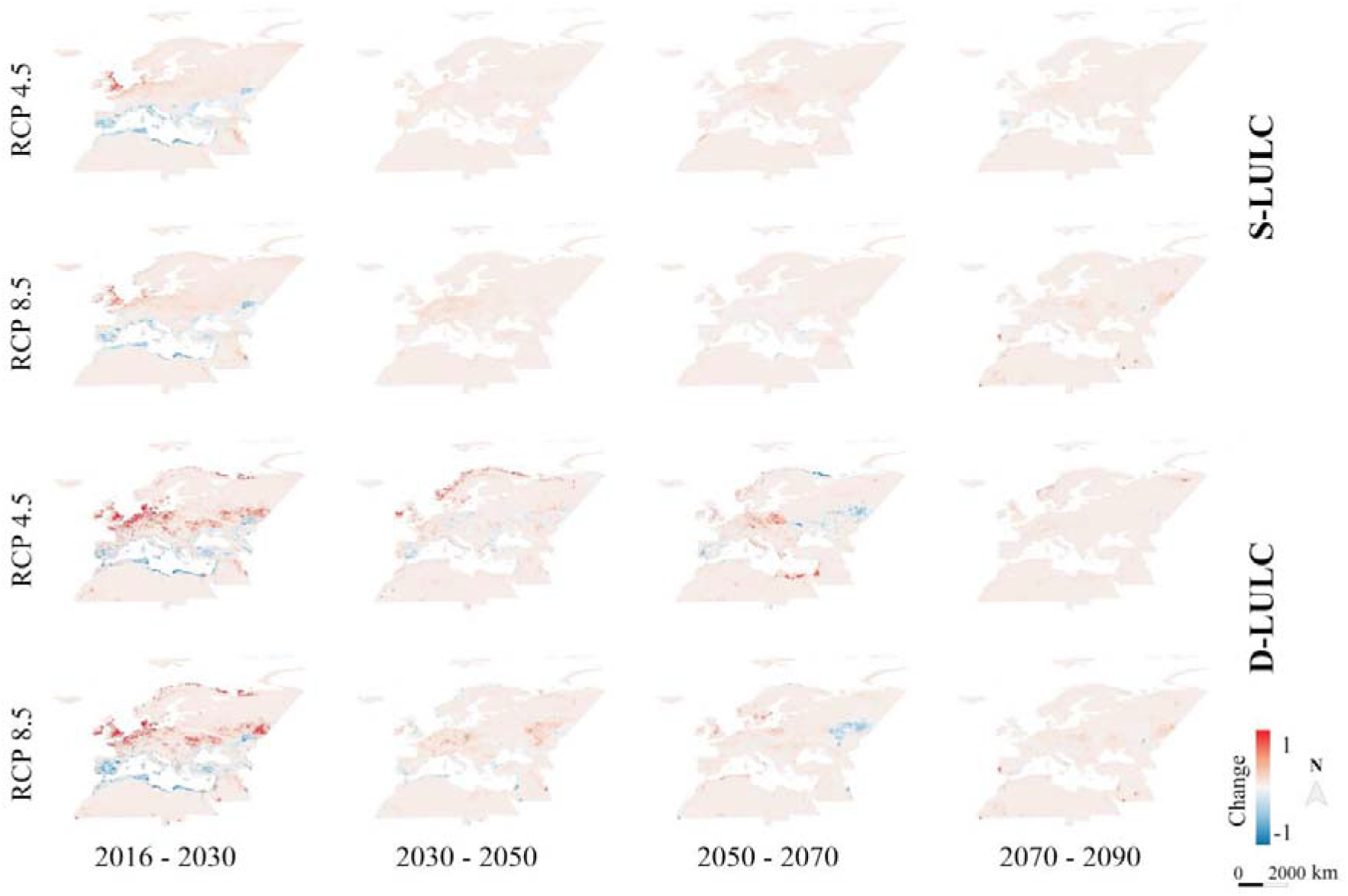
Cell-wise change of breeding habitat suitability for the Eurasian stone-curlew under four time intervals (2016-2030, 2030-2050, 2050-2070, 2070-2090) and two Representative Concentration Pathways (RCP 4.5, RCP 8.5). S-LULC = Static Land-Use/Land-Cover; D-LULC = Dynamic Land-Use/Land-Cover.

### Change of mean habitat suitability

The mean breeding habitat suitability considering a static LULC is predicted to remain stable in all time intervals and for most GCMs (Fig. S1.6A and S1.6B). However, when comparing the predictions for different GCMs, the differences were quite marked. The RCP had also a significant effect, as the percentages under RCP 8.5 were remarkably higher (Fig. S1.6C).

### Critical areas for conservation

France, South-Eastern UK, Northern Italy and Israel were the regions with the highest proportion of conservation critical areas (Fig. 4). The percentage of currently suitable areas predicted to remain suitable was relatively low (27.47 %).

**Figure 4.**
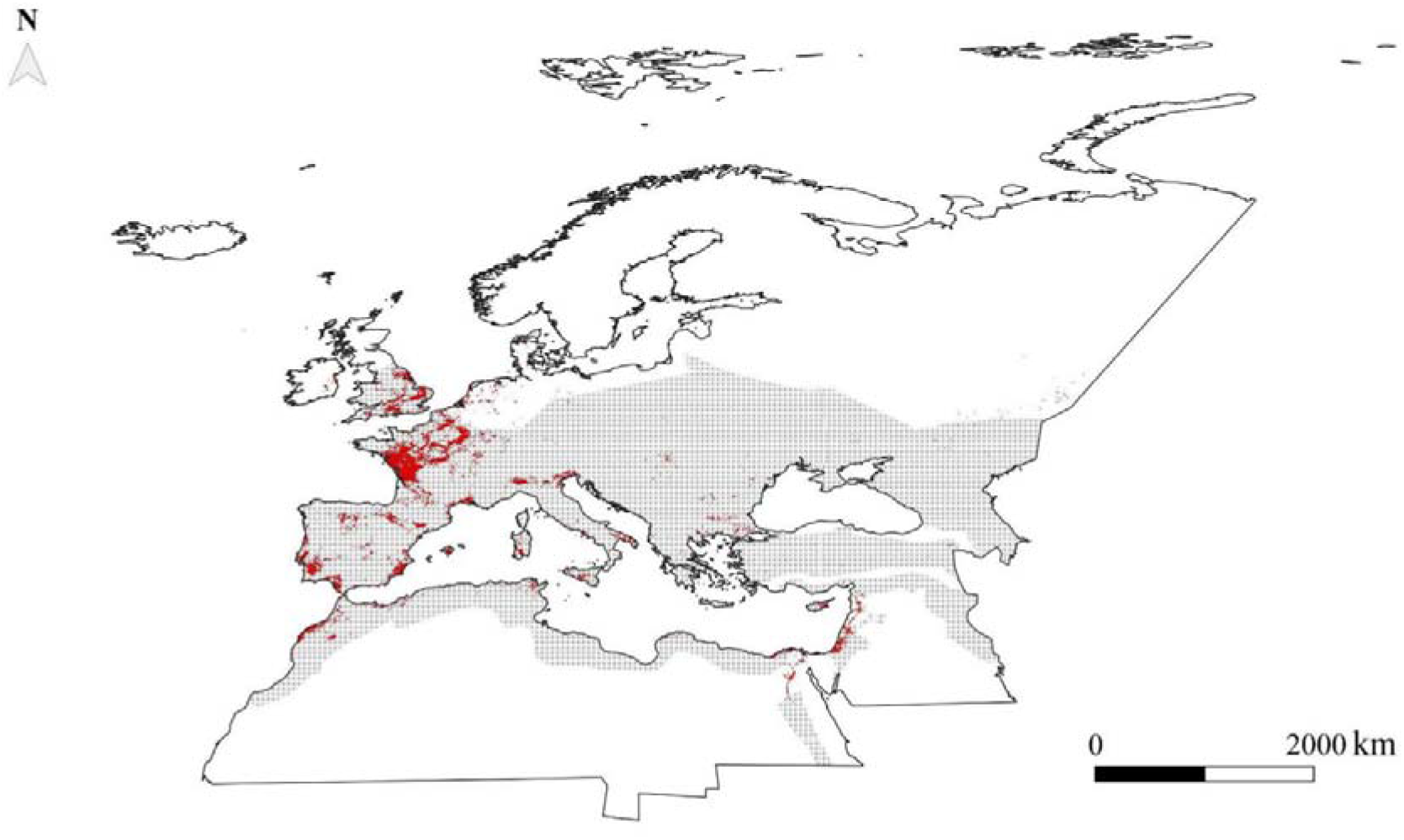
Critical areas for the conservation of the Eurasian stone-curlew. Currently suitable areas predicted to remain suitable under all scenarios of breeding habitat suitability for the species are highlighted in red. Shaded areas represent the species’ range in the Western Palearctic, with the exclusion of the Macaronesian archipelago.

## DISCUSSION

Relying on 1628 occurrences of the stone-curlew, we developed a robust species distribution modelling framework for the species, based on a Maxent model built with year-specific variables and calibrated using a kernel density approach for the pseudo-absences. Our model predicted significant short-term changes in the spatial distribution of suitable areas for the species and allowed the identification of areas critical for its conservation under global change. Furthermore, our results suggested that the distribution of the stone-curlew might be affected by short-term conditions, rather than long-term ones, and confirmed that broad areas of the Western Palearctic are likely to be suitable for the species in the future.

The northern shift of suitable areas predicted by our model is consistent with our hypothesis, with the projection for the species (Huntley *et al*., 2007), and with projections for other pseudo-steppic species (e.g. the Great Bustard *Otis tarda* and Little Bustard *Tetrax tetrax* in Estrada *et al*., 2016 and the European Roller *Coracias garrulus* in Kiss *et al*., 2020). The loss of suitable areas in the Mediterranean region is not surprising as the Mediterranean biogeographical region represents a hotspot of global change, and extensive biodiversity losses have been predicted (Sala *et al*., 2000).

Even if strong variations between GCMs and RCPs in the expected trajectories of mean breeding habitat suitability for the stone-curlew may hinder sound conclusions, the predicted gains and losses of mean habitat suitability were balanced, suggesting a stable mean habitat suitability for the species through time. This agrees with our expectation that mean habitat suitability for the species would remain stable or increase, as global warming is milder in warm temperate areas (IPCC, 2013) and might therefore have a limited impact on mean habitat suitability. Dispersal ability, biotic interactions and the carrying capacity of suitable habitats might determine whether a stable mean habitat suitability translates into a stable population size (Bateman *et al*., 2013; Holloway, Miller, & Gillings, 2016).

Habitat suitability has been linked to population size and trends in birds (Green *et al*., 2008; Stiels *et al*., 2021). We showed that large areas of the study region are predicted as suitable in the future, and that a stable mean habitat suitability is likely. Hence, according to our forecasts, the species may be able to maintain viable populations in the Western Palearctic. However, given the relatively low percentage of areas suitable under both current and future conditions and the predicted shift of suitable areas, the species might need to track its niche in the future. Evidence of the ability of animals to track their niche is contrasting (Devictor *et al*., 2008; Chen *et al*., 2011) and many terrestrial organisms have been shown to shift their distribution at a sufficient pace to track recent temperature changes (Chen *et al*., 2011).

Species affected by global change may also persist under unfavourable conditions being phenotypically plastic and becoming locally adapted (Valladares *et al*., 2014). Moreover, biological interactions and movement/dispersal constraints might prevent them from colonizing newly suitable areas (Bateman *et al*., 2013; Holloway, Miller, & Gillings, 2016). Our results point toward a critical use of the target-group pseudo-absence method, and we recommend its explicit testing especially when using multi-source citizen science datasets. Our results did not evidence a better performance for ensemble methods, contrary to other studies (e.g. Marmion *et al*., 2009), and support a tailor-made choice of the modelling method. Dynamic models performed better than static ones (Table 3) as expected, supporting previous works explicitly testing the two approaches (e.g. Reside *et al*., 2010; Milanesi *et al*., 2020a). Due to its minimal contribution to the final model, the ‘prey suitability’ variable (Fig. S1.1) probably failed to describe the real patterns of habitat suitability for the stone-curlew’s preys (Table 1). This might depend on the exclusion of relevant prey items, as the species exploits a wide spectrum of food items (Vaughan & Vaughan-Jennings, 2005) that is difficult to model through SDMs. We nevertheless highlight the importance of considering biotic predictors though suggest including the entire trophic web of a species.

Correlative species distribution models assume an equilibrium condition between the species and the environment (Araújo & Pearson, 2005). Areas that have been abandoned by the stone-curlew during the last decades of the past century can be recolonized, as happened in the UK following the targeted conservation efforts of the LIFE11INF/UK000418 ‘Securing the future of the stone-curlew throughout its range in the UK’. This suggests that suitable climate and LULC conditions at the relatively coarse scale of our study exist in those areas. Therefore, the SDMs’ equilibrium assumption for this species might be violated and future studies could explicitly account for non-equilibrium conditions (Václavík & Meentemeyer, 2012). SDMs also assume the evolutionary conservatism of the species’ ecological niche through time (Peterson, Soberón, & Sanchez-Cordero, 1999; Barbet-Massin *et al*., 2011), a condition that is difficult to prove and deserves specific attention. Thanks to constant improvements in climate and LULC scenarios, in the future our workflow could be applied to a wider spectrum of scenarios, to better represent all the components of uncertainty (Thuiller *et al*., 2019). However, the two RCPs employed in our study can provide valuable insights on the effect of global change on habitat suitability, as demonstrated by their extensive use (e.g. Kiss *et al*., 2020; Stiels *et al*., 2021). We also limited our study to the breeding period: understanding the drivers and changes of habitat suitability for the stone-curlew across migratory and wintering grounds might provide further insights for its conservation.

The stone-curlew is philopatric in the breeding areas (Green, 1990) and this might delay habitat-tracking under environmental change. Considering this latter scenario, critical areas might act as a stronghold for the species, and anticipatory conservation efforts should primarily focus on these areas (Thuiller *et al*., 2019). Furthermore, ensuring high connectivity conditions for the species in areas predicted to become unsuitable might contribute to maintain viable populations and facilitate niche tracking (Heller & Zavaleta, 2009). Indeed, habitat fragmentation contributed to the species’ decline in the ‘90s (Tucker & Evans, 1997). Finally, enhanced monitoring efforts in areas predicted to become suitable might increase early-detection probability and mitigate the negative effects of biotic interactions. Our model evidenced the importance of two LULC classes that are heavily affected by farming and harvesting activities (i.e. non-irrigated agricultural areas and grasslands). Precisely, the most critical areas and the areas predicted to become suitable for the stone-curlew are in intensively cultivated areas, corresponding to the ‘core of EU continental agriculture’ (D’Amico *et al*., 2013), England and the Po Plain. Furthermore, in France, the country hosting the largest portion of critical areas, over 60% of breeding pairs are found in the Central/Western region within arable crops (Malvaud, 1996; Issa & Muller, 2015). In these ecosystems, *ad hoc* management interventions on a local scale can be effectively used to favour stone-curlew’s presence (Hawkes *et al*., 2021).

In conclusion, our study refines current knowledge on the effects of global change for the stone-curlew, providing predictions of breeding habitat suitability for the species at high temporal and spatial resolutions, and conservation planners might benefit from our results by incorporating indications on the most relevant conservation areas in the development of action plans for the species.

## ACKNOWLEDGMENTS

We thank Chris Vernon, Diego Rubolini, Luca Forneris and Min Chen for technical help, Danae Portolou and Nikos Tsiopelas for providing aggregated occurrence data. We also thank Saverio Gatto for granting us the permission to use his stone-curlew’s photograph.

## DATA ACCESSIBILITY STATEMENT

Presence data used to develop species distribution models for the Eurasian stone-curlew are available at: https://doi.org/10.6084/m9.figshare.16727296.v1

Longitude, latitude and year are reported for each observation. Data from the British Trust for Ornithology are omitted in agreement with the data access policy. Data from ornitho.it are accompanied by a citation, in compliance with the site’s rules.

## List of Appendices

**Appendix S1. Overview, Data, Model, Assessment and Prediction (ODMAP) protocol (Zurell *et al*. 2020).**

