## Appendix S1 for "Projected dynamics of breeding habitat suitability for a steppe-land bird warrant anticipatory conservation actions"

#### – ODMAP Protocol –

Andrea Simoncini, Samuele Ramellini, Alexis Martineau, Alessandro Massolo, Dimitri Giunchi

Date: to be updated upon acceptance

---

#### Overview

##### Authorship

A. Simoncini and S. Ramellini should be considered as joint first authors.

Author contribution: AS, SR, AMas, DG (Conceptualization); AS, SR, AMar (Data mining); AS, SR (Analysis); AS, SR (First draft); AS, SR, AMas, DG (Manuscript revision). All the authors approved the final version.

Contact:

Study link: to be updated upon acceptance.

##### Model objective

Model objective: predict current and future breeding habitat suitability for a steppe-land bird, quantify its dynamics and identify critical conservation areas.

Target output: maps of current (2016) and future (2030, 2050, 2070, 2090) breeding habitat suitability; map of cell-wise change of breeding habitat suitability between consecutive time periods; description of mean change of breeding habitat suitability between consecutive time periods; map of areas critical for the conservation of the focal taxon.

##### Focal Taxon

Focal Taxon: Eurasian stone-curlew (*Burhinus oedicnemus*, hereafter stone-curlew).

#### Location

<u>Location:</u> Western Palearctic (*sensu* Snow & Perrins, 1998).

#### Scale of Analysis

<u>Spatial extent:</u> -24.53991, 80.43719, 17.49258, 81.85871 (xmin, xmax, ymin, ymax).

<u>Spatial resolution:</u> 0.05° resolution (i.e. ~ 5.5x5.5 km grid cells).

<u>Temporal extent:</u> 1992-2016 (prediction: 2030, 2050, 2070, 2090).

<u>Temporal resolution:</u> one year.

<u>Boundary:</u> natural (Biogeographic region).

#### Biodiversity data

<u>Observation type:</u> citizen science, field survey, standardised monitoring data.

<u>Response data type:</u> presence-only.

#### Predictors

<u>Predictor types:</u> predictors were represented by climatic, Land-Use/Land-Cover (LULC), edaphic and topographic variables. We compared static and dynamic SDMs (*sensu* Milanesi, Della Rocca, & Robinson, 2020). Static SDMs relate temporally dynamic occurrences to static predictor variables (e.g. 30-year climate averages), yielding potentially biased estimations of species-environment relationships (Milanesi *et al.,* 2020). Instead, dynamic SDMs combine species records with time-specific raster values and provide better results (Reside *et al*., 2010; Milanesi *et al.,* 2020).

#### Hypotheses

<u>Hypotheses:</u> we hypothesized that stone-curlew's range would shift northwards and that large areas of the Western Palearctic will be suitable for the species in the future, as reported for other steppe-land birds (e.g. the European bustards *Otis tarda*/*Tetrax tetrax* in Estrada *et al*., 2016; the roller *Coracias garrulus* in Kiss *et al*., 2020). We also hypothesized that dynamic models would perform better than static ones (Reside *et al*., 2010; Milanesi *et al.,* 2020).

#### Assumptions

Model assumptions: As for most studies employing correlative species distribution models, we assumed an equilibrium between the species and the environment (Araújo & Pearson, 2005) and the evolutionary conservatism of the ecological niche (Peterson, Soberón, & Sanchez-Cordero, 1999).

#### Algorithms

Modelling techniques: as no 'silver bullet' technique exists in species distribution modelling (Qiao, Soberón, & Peterson, 2015), we tested seven single algorithms (ann, cta, gam, gbm, glm, maxent, randomForest; see Table S1.1). We also combined the single techniques to produce two ensembles, according to the mean and the weighted average (Marmion *et al.*, 2009) and contrasted them with the single algorithms during the model-selection phase.

**Table S1.1** List of candidate species distribution modelling algorithms used to describe breeding habitat suitability for the Eurasian stone-curlew in the model-selection phase. For each algorithm the key reference is reported.

| Algorithm | Abbreviation | Key reference |
|---|---|---|
| Artificial Neural Networks | ANN | Ripley, 2007 |
| Classification Tree Analysis | CTA | Breiman, 1984 |
| Generalized Additive Model | GAM | Hastie & Tibshirani, 2004 |
| Generalized Boosting Model | GBM | Elith, Leathwick, & Hastie, 2008 |
| Generalized Linear Model | GLM | McCullagh, 1984 |
| Maximum Entropy | Maxent | Phillips, Anderson, & Schapire, 2006 |
| Random Forest | RF | Breiman, 2001 |

Model complexity: to avoid exceedingly complex models (with a high overfitting), we computed the difference between the AUC for the training dataset and the test dataset ($AUC_{diff}$). High values indicated overly complex models (Radosavljevic & Anderson, 2014).

#### Workflow

Model workflow: we first tested two pseudo-absence methods, to develop a pseudo-absence dataset that effectively canceled out the spatial bias in the presence dataset. We then tested four modelling strategies:

(i) single static SDMs: seven algorithms were developed (see Table S1.1 for a list of the employed techniques), using static variables;

(ii) ensemble SDMs: the single static SDMs were combined according to the mean and the weighted average (Marmion *et al*., 2009). In the weighted average method, the AUCtest was used to weight model contribution to the final prediction. Static variables were employed;

(iii) single dynamic SDMs: seven algorithms (Table S1.1) were employed, using dynamic variables;

(iv) ensemble dynamic SDMs: the single dynamic SDMs were combined according to the mean and the weighted average, using dynamic variables.
We then projected the best model to derive maps of breeding habitat suitability for the stone-curlew under a set of future scenarios. Using these projections, we computed the cell-wise and mean change of breeding habitat suitability between consecutive time intervals. Finally, we identified the areas predicted as suitable under both current and future suitability scenarios, and therefore critical for the species 'conservation (Estrada *et al*., 2016; Thuiller *et al*., 2019).

#### Software

Software: R version 4.0.4 (2021-02-15).

Code availability: available from the corresponding authors upon reasonable request.

Data availability: https://doi.org/10.6084/m9.figshare.16727296.v1

### Data

#### Biodiversity data

Taxon names: Eurasian stone-curlew (*Burhinus oedicnemus*).

Taxonomic reference system: Integrated Taxonomic Information System (ITIS).

Ecological level: species.

Data sources: eBird (eBird.org); Global Biodiversity Information System (GBIF.org); ornitho.it; ornitho.cat; xeno-canto.org; British Trust for Ornithology (obtained from the coordinators of the database); Ornithotopos database and the second European Breeding Bird Atlas (EBBA) 2x2 km surveys in Greece (in formal agreement with the Hellenic Ornithological Society), data for the Deux-Sèvres department from the Nature79 database provided by Alexis Martineau, and nest points from Northern Italy (collected by Dimitri Giunchi).

Sampling design: citizen Science and uniform counts (EBBA counts in Greece).

Sample size: 1628 presences.

Absence data: not used.

Background data: we obtained a set of pseudo-absences (used also as background data for Maxent models) generating a large number of pseudo-absences and weighting their spatial distribution according to a fixed kernel density estimated on all the presence points, by means of the 'adehabitatHR' (v. 0.4.18) R package (Calenge, 2006). We used the *ad hoc* method for the estimation of the smoothing parameter (Worton, 1989). The procedure was independently repeated on the occurrences separated by year for the period 1992-2016. If a cell contained more than one record, the chronologically oldest was retained in the dataset. Finally, we removed pseudo-absences outside the study area and randomly sampled them to obtain a 0.1 prevalence (Barbet-Massin, Thuiller, & Jiguet, 2012).

#### Data partitioning

We evaluated models based on the block cross-validation procedure implemented in the 'ENMeval' (v. 0.3.1) R package (Muscarella *et al*., 2014). In this procedure, data are partitioned into four geographically-independent bins; k models are then produced, with k-1 bins used for training and the remaining ones used for testing. We selected k = 4 (i.e. a four-fold spatial block cross-validation).

#### Predictor variables

<u>Predictor variables:</u> we used an expert-based approach for model calibration (Santini *et al*., 2021) to define 17 variables representing climate, topography, soil composition and Land-Use/Land-Cover (LULC).

Climate is a major determinant of stone-curlew's distribution at the broad scale (Vaughan & Vaughan-Jennings, 2005), as the species selects warm and dry areas to reproduce (Green, Tyler, & Bowden, 2000; Vaughan & Vaughan-Jennings, 2005; Keller *et al*., 2020). We therefore included a set of variables describing temperature (annual temperature, temperature of the warmest quarter and temperature seasonality) and precipitation (annual precipitation, precipitation of the warmest quarter and precipitation seasonality).

LULC change affects the distribution and abundance of the stone-curlew (Burfield, 2005; Onrubia & Andrés, 2005). Agricultural areas with low vegetation density are exploited by stone-curlews for breeding and foraging (Vaughan & Vaughan-Jennings, 2005; Caccamo *et al*., 2011). Arid and steppic grasslands are among the elective habitats for the species (Green *et al.,* 2000; Vaughan & Vaughan-Jennings, 2005; Hume & Kirwan, 2013; Teyar *et al*., 2020), as well as low shrubs, often mixed with grass and bare ground (Vaughan & Vaughan-Jennings, 2005; Traba *et al*., 2013). On the contrary, trees are often counter-selected (Madroño, González, & Atienza, 2004; Vaughan & Vaughan-Jennings, 2005). Breeding attempts in urbanized areas are increasingly reported (Cutini, Campedelli, & Tellini Florenzano, 2006; Biondi *et al*., 2015; Giovacchini *et al*., 2017). We derived the percentage cover for the following classes: i) agricultural (non-irrigated), ii) agricultural (irrigated), iii) grass, iv) shrub, vi) tree, vii) bare and viii) urban. These classes adequately represent the main LULC drivers of habitat suitability for the species previously described.

Grounds with a slope lower than 10° are preferred by breeding stone-curlews (Vaughan & Vaughan-Jennings, 2005). Therefore, we included a variable representing slope.

A preference of the stone-curlew for areas close to inland water has been reported (Vaughan & Vaughan-Jennings, 2005; Caccamo *et al*., 2011). We therefore produced a variable representing the distance of the centroid of each cell to the nearest source of inland water.

The stone-curlew breeds on calcareous soils in many areas of its range (e.g. in Italy, France and England; (Vaughan & Vaughan-Jennings, 2005; Biondi *et al*., 2015). These soils are typically well-drained and dry, conditions that are highly conducive to the species 'presence (Cramp & Simmons, 1983; Vaughan & Vaughan-Jennings, 2005; Gaget *et al*., 2019). We therefore produced a raster variable describing the presence of karst areas. In this raster, a pixel had a value of 0 if it was outside a karst patch, and the area of the corresponding karst patch if it was within a patch.

Biological interactions contribute to shape species responses to global change (Araújo & Luoto, 2007; Wisz *et al*., 2013). We therefore included a proxy of prey availability among the explanatory variables. We produced a static Maxent model based on prey occurrence data extracted from the GBIF repository. We downloaded all the available data for Lumbricidae, Acrididae, Scarabaeidae, Carabidae and Curculionidae, i.e. the invertebrate families identified as stone-curlew's prey by at least four papers in the most extensive review of the species 'feeding habits (Vaughan & Vaughan-Jennings, 2005; Giovacchini *et al*., 2017). After removing duplicates, to reduce computation time we randomly sampled 1000 occurrences for each major taxonomic group (i.e. Oligochaeta, Orthoptera, Coleoptera), resulting in 3000 presences. A kernel-density approach was used to define pseudo-absences (see *Pseudo-absence selection* in the main text). A four-fold spatial block cross-validation strategy was then used to define training and test datasets. Environmental conditions were represented by static variables with a VIF < 3 (Zuur, Ieno, & Elphick, 2010), resulting in the exclusion of mean annual temperature, mean annual precipitation, precipitation seasonality and percentage cover of bare land. The model produced with the first cross-validation run had the best performance and was selected for subsequent analyses (AUCtest = 0.684, CBI = 0.945). We then projected the model on the static variables to represent mean prey suitability (Figure S1.1a) and produced year-specific projections for the period 1992-2016 (Figure S1.1b, Figure S1.1c) to use as input for the dynamic prey suitability variable, obtained with the standard procedure adopted for dynamic variables. The same model was also projected on the future variables to forecast future prey suitability (Figure S1.1d).

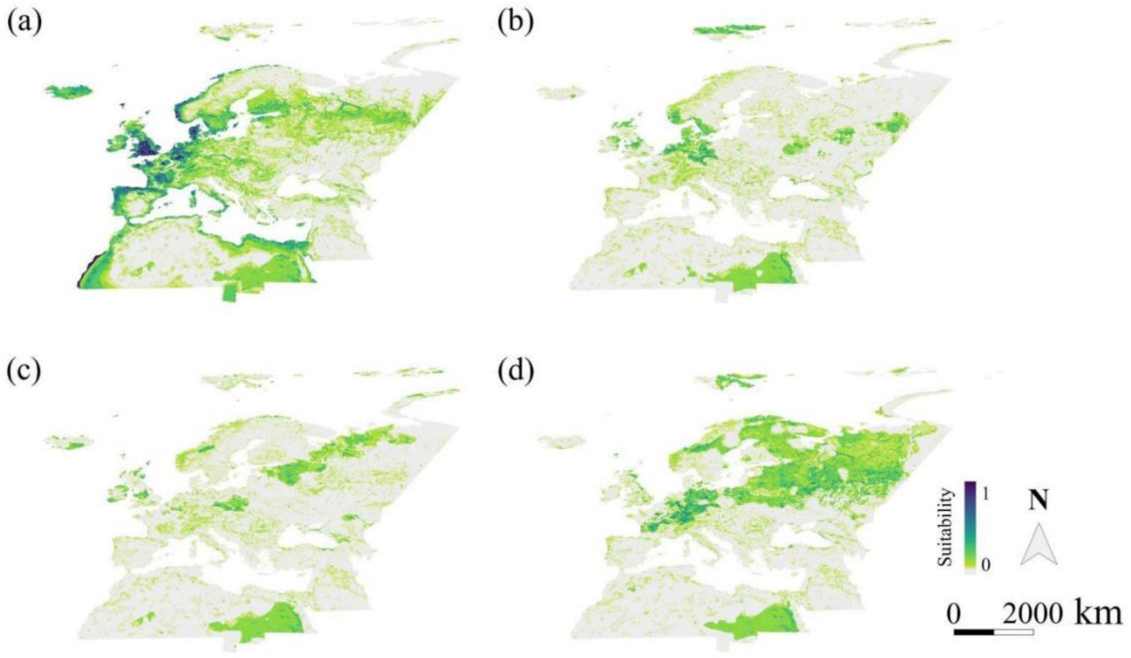


**Figure S1.1** Example outputs from the prey suitability species distribution model, describing habitat suitability for Eurasian stone-curlew's preys. (a) Model projection on the static variables, (b) model projection on the 2016 variables, (c) model projection on the 1992 variables, (d) model projection on the 2050 variables (General Circulation Model ACCESS1-3, Representative Concentration Pathway 8.5).

Data sources: climate variables were retrieved from the CHELSAcruts database (Karger & Zimmermann, 2018). We used the global land cover developed by the European Space Agency

(European Space Agency, 2019) to describe LULC. To account for the effect of topography we derived a slope variable from the 'SEDAC Altimeter Corrected Elevations (ACE2)' (version 2.0). To describe distance from water, we used the 'World Water Bodies' (considering only inland features) and the 'World Linear Water' shapefiles available on ArcGIS Hub (www.hub.arcgis.com). We used the 'World Map of Carbonate Rock Outcrops' (version 3.0) to produce a raster representing karst patches (Mammola *et al.*, 2019).

Spatial extent: all variables were cropped to the Western Palearctic.

Spatial resolution: all variables were resampled to the 0.05° resolution.

Coordinate reference system: World Geodetic System 1984.

Temporal extent: static variables were represented by averaged values for the years 1973-2013 (climatic and prey suitability variables) and 1992-2016 (LULC variables). For dynamic SDMs, we associated occurrence/pseudo-absence pixels with year-specific values for a given variable.

##### Transfer data

Data sources: CHELSA CMIP-5 timeseries (Karger *et al.*, 2017), global LULC scenarios from Chen *et al.* 2020.

Spatial extent: Western Palearctic.

Spatial resolution: 0.05° (~ 5.5x5.5 km).

Temporal extent: 2016 (considered as current), 2030, 2050, 2070, 2090.

Models and scenarios: we initially used a static LULC (i.e. the 2016 LULC variables) in future predictions to constrain climatic outputs (Pang, De Alban, & Webb, 2021). We then performed an additional analysis projecting models calibrated with the same historical LULC database (ESA - European Space Agency 2019) on scenarios from the future dynamic LULC database (Chen *et al.*, 2020). In static LULC projections, the four models selected were ACCESS1-3 (Dix *et al.*, 2013), CESM1-BGC (Lindsay *et al.*, 2014), CMCC-CM (Scoccimarro *et al.*, 2011) and MIROC5 (Watanabe *et al.*, 2010). We considered two emission scenarios corresponding to the IPCC's Representative Concentration Pathways 4.5 and 8.5 (RCP 4.5 and RCP 8.5). For each RCP scenario and time interval we developed 4 projections (one per each GCM) averaging them in a final projection representing habitat suitability for the species. In dynamic LULC projections, we developed a single projection for each RCP scenario and time interval, under the Shared Socioeconomic Pathway (SSP) 5 and the MIROC circulation model.

Quantification of Novelty: novelty was quantified according to the environmental overlap mask (Zurell, Elith, & Schröder, 2012), considering both strict and combinational extrapolation.

### Model

#### Multicollinearity

We calculated the Variance Inflation Factor (VIF) to address multicollinearity between predictors. We used the vifstep function in the 'usdm' R package (v 1.1-18; Naimi, 2013), using a threshold of 3 (Zuur *et al.*, 2010). We report the initial and final VIF values for static and dynamic variables in Table 1 (see main text).

#### Model settings

Default 'biomod2' (v. 3.3-7; Thuiller *et al.*, 2016) settings were used to develop individual SDM algorithms.

#### Model estimates

To establish the contribution of each variable in a model, we used the model independent approach (Thuiller *et al*., 2009). We calculated Pearson's correlation (r) between the fitted values from the original dataset and those where the focal variable had been randomly permutated. A high correlation coefficient defines a scarcely important variable. We expressed the variable importance as percentage.

#### Analysis and Correction of non-independence

We didn't account explicitly for spatial autocorrelation in this study. However, we buffered the negative effects of possible non-independence by spatially thinning presence data with the 'spThin' R package (Aiello-Lammens *et al*., 2015), using a distance between records of 5.5 km.

#### Threshold selection

We used the threshold that maximizes the sum of specificity and sensitivity, a sound thresholding method (Liu, Newell, & White, 2016).

### Assessment

#### Performance statistics

Performance on training data: To assess model performance on the training data we used the Area Under the Receiver-Operating Characteristic (ROC) Curve (AUC, Fielding & Bell, 1997), computed on the training dataset (AUCtrain).

Performance on test data: To assess model performance on the test data we used the AUC computed on the test dataset (AUCtest) and the Continuous Boyce Index (CBI, Boyce *et al.*, 2002). The difference between AUCtrain and AUCtest ($AUC_{diff}$) was also computed to quantify overfitting (Radosavljevic & Anderson, 2014).

#### Plausibility check

Response shapes: We produced the response curves for each model using the evaluation strip method (Elith *et al.*, 2005). Response curves for the four variables with the highest percentage contribution in each model were evaluated through a subjective score, defined Plausibility of Response curve (PRC). The PRC was assigned by two authors (AS, SR) based on the agreement between current knowledge on the species' ecological preferences and the model's selected response curves.

Expert judgement: Spatial projections were evaluated through a subjective score (Perennes *et al.*, 2021), defined Quality of Spatial Prediction (QSP). The QSP was assigned by two authors (AS, SR) based on expert knowledge and comparing qualitatively the spatial output of each model with the species' distribution in the EBBA1 and EBBA 2 (Hagemeijer & Blair, 1997; Keller *et al.*, 2020) and with the modelled distribution in the EBBA 2 (Keller *et al.*, 2020). Both QSP and PRC scores had three possible values (zero, one, two), with higher values indicating a better performance.

### Prediction

#### Prediction output

Spatial outputs from models built with the two pseudo-absence methods are presented in Figure S1.2, whereas the outputs of static and dynamic models are reported in Figures S1.3 and S1.4. Future projections from the final model (dynamic Maxent) are found in Figure S1.5, whereas the cell-wise and mean change of breeding habitat suitability are respectively described in Figure 4 (main text) and Figure S1.6.

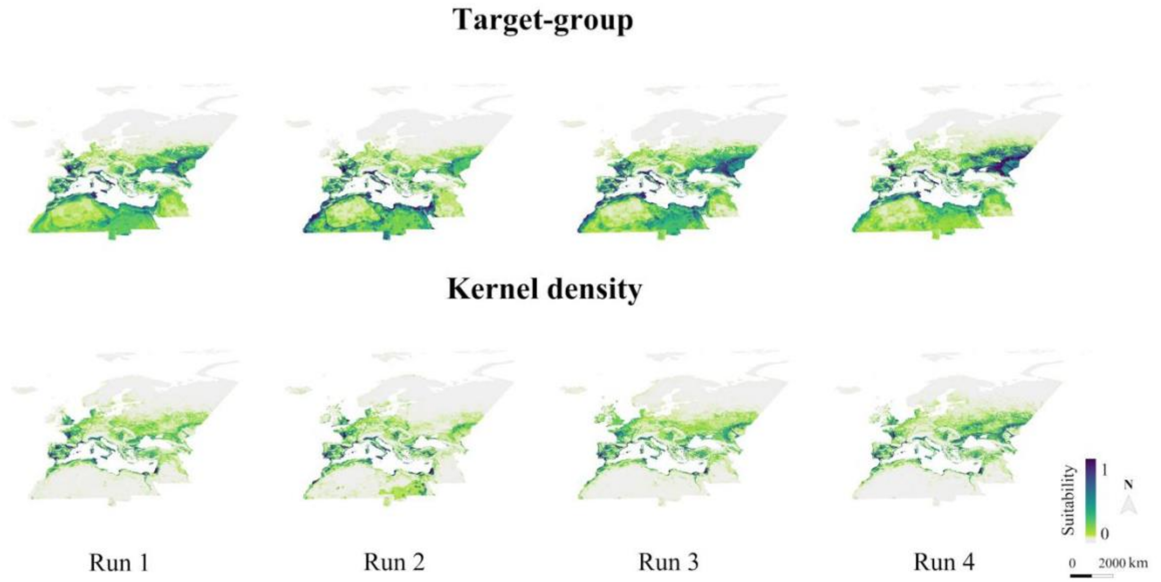


**Figure S1.2** Maps of breeding habitat suitability for the Eurasian stone-curlew under current conditions according to two pseudo-absence methods (‘Target-group ’and ‘Kernel density’). Run 1-4 represent the four runs developed in a four-fold spatial block cross-validation strategy.

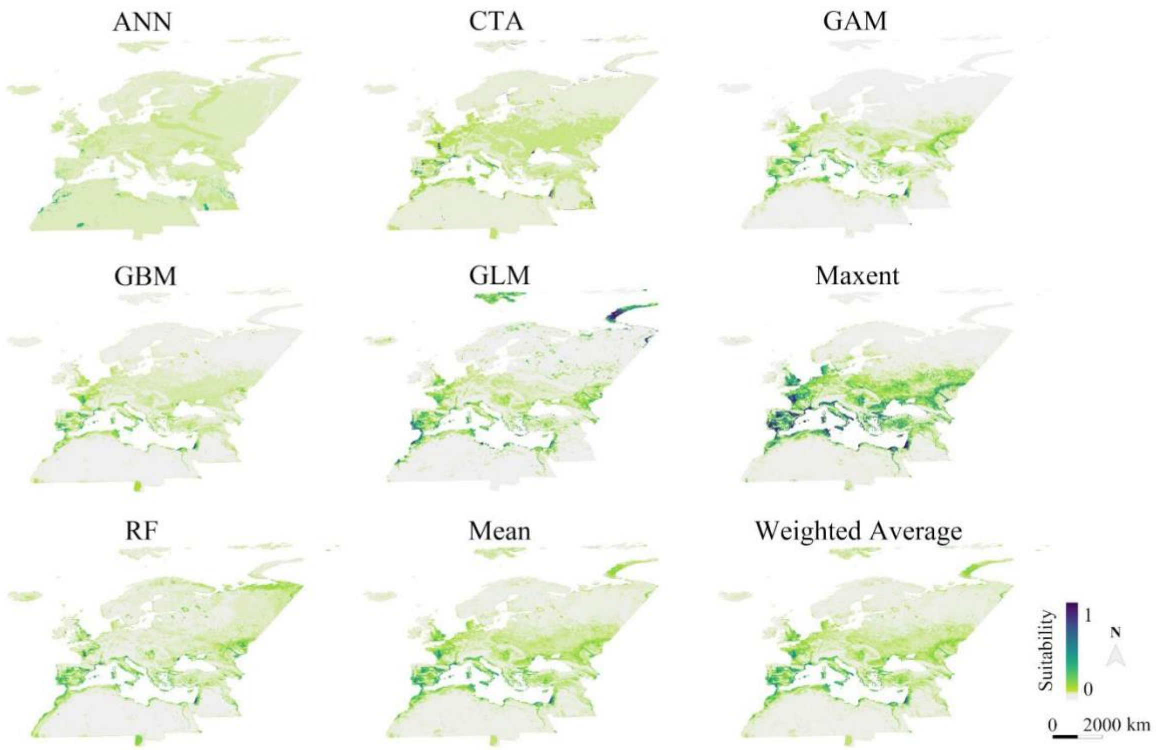


**Figure S1.3** Maps of breeding habitat suitability for the Eurasian stone-curlew according to static species distribution models under current conditions. Projections of seven single algorithms (ANN = Artificial Neural Networks, CTA = Classification Tree Analysis, GAM = Generalized Additive Model, GBM = Generalized Boosting Model, GLM = Generalized Linear Model, Maxent = Maximum Entropy, RF = Random Forest) and two ensemble methods (mean and weighted average) are reported.

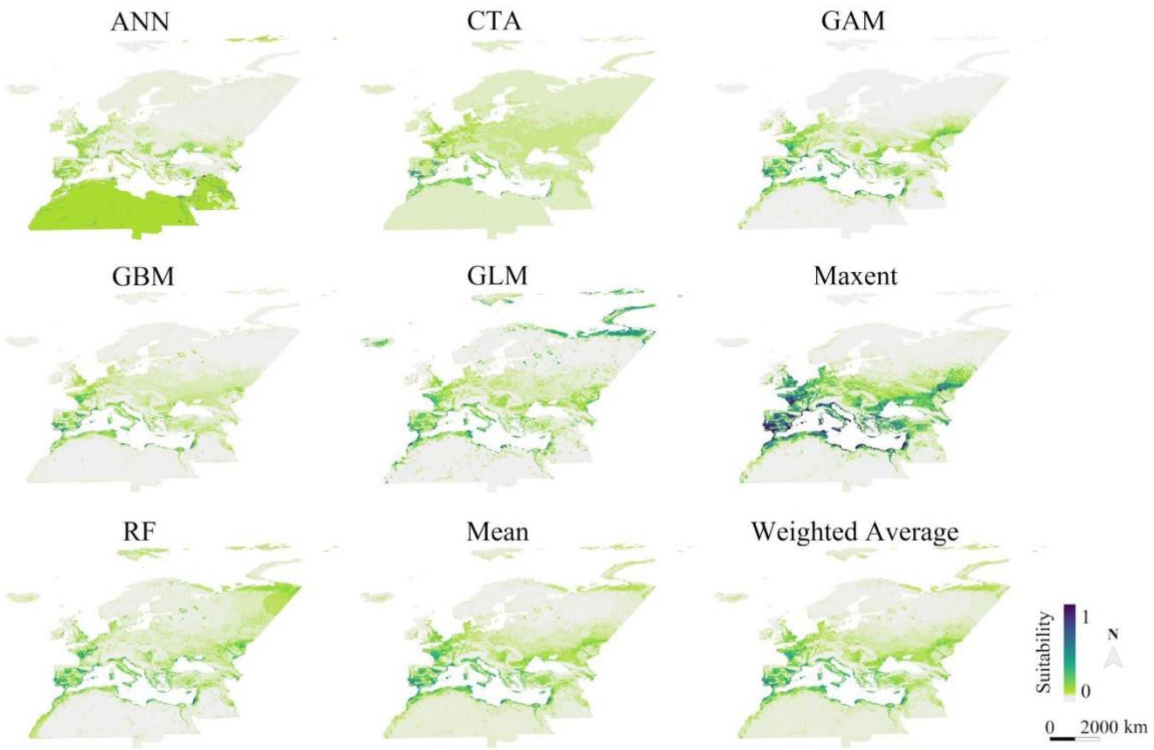


**Figure S1.4** Maps of breeding habitat suitability for the Eurasian stone-curlew according to dynamic species distribution models under current conditions. Projections of seven single algorithms (ANN = Artificial Neural Networks, CTA = Classification Tree Analysis, GAM = Generalized Additive Model, GBM = Generalized Boosting Model, GLM = Generalized Linear Model, Maxent = Maximum Entropy, RF = Random Forest) and two ensemble methods (mean and weighted Average) are reported.

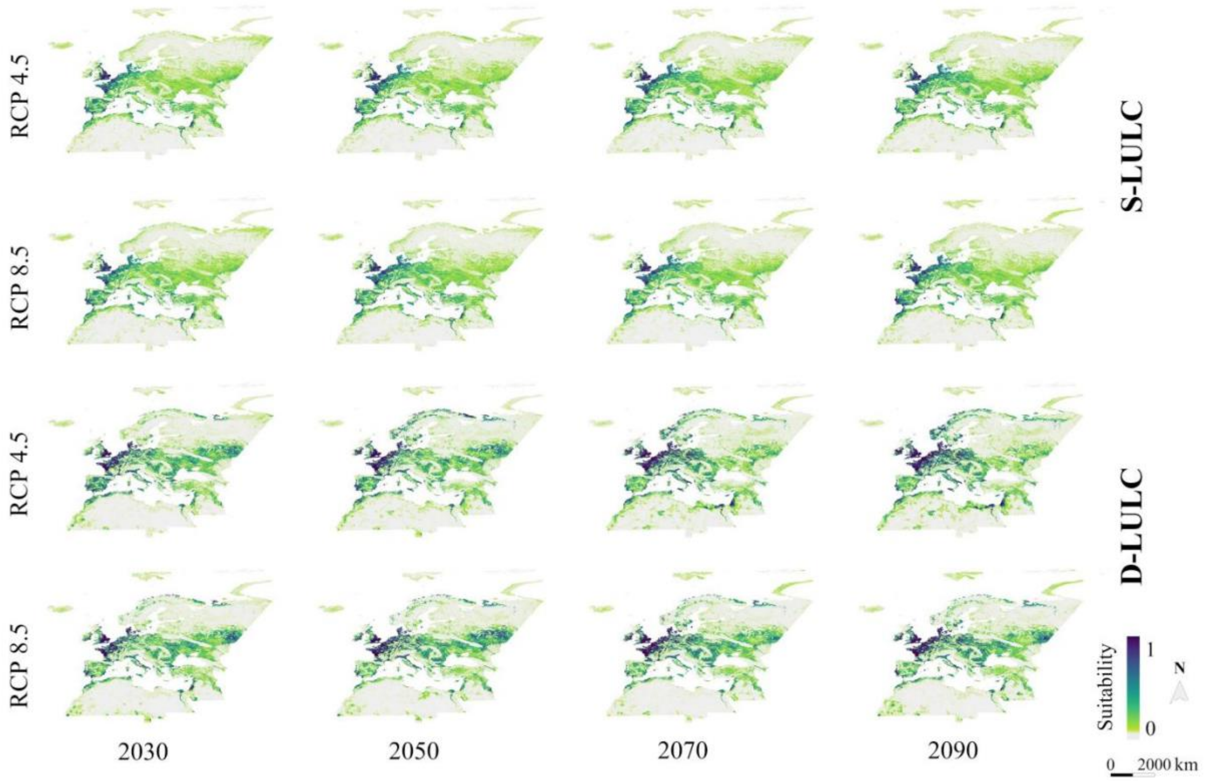


**Figure S1.5** Projections of breeding habitat suitability for the Eurasian stone-curlew under two Representative Concentration Pathways (RCP 4.5, RCP 8.5) and four time periods. In Static Land-Use/Land-Cover (S-LULC) projections, current LULC conditions were used; in Dynamic Land-Use/Land-Cover (D-LULC) projections, LULC conditions changed according to the Shared Socio-economic Pathway 5 (SSP 5). For S-LULC projections, each map represents the mean of projections under four General Circulation Models; for D-LULC projections, only the MIROC circulation model was used.

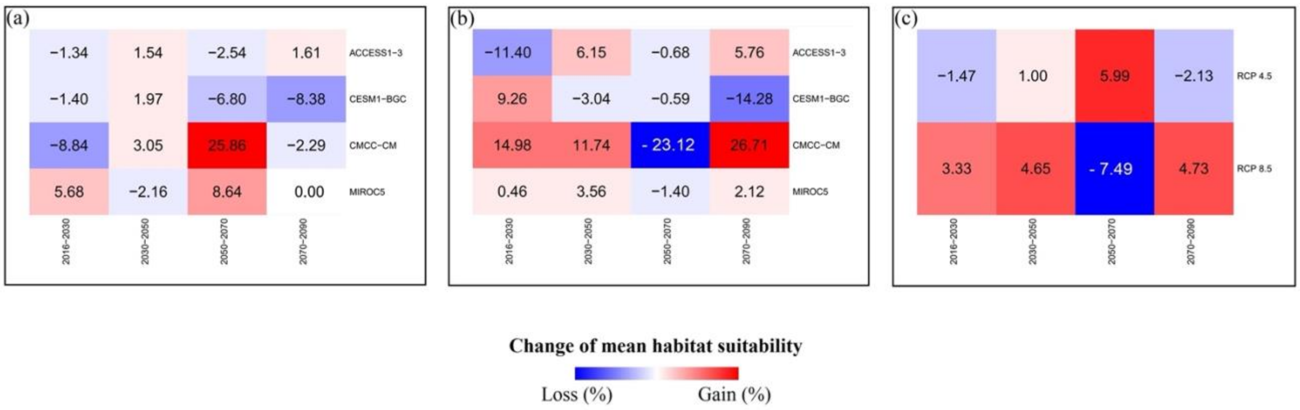

**Figure S1.6** Change of mean breeding habitat suitability for the Eurasian stone-curlew under four time intervals (2016-2030, 2030-2050, 2050-2070, 2070-2090), according to four General Circulation Models (GCMs: ACCESS1-3, CESM1-BGC, CMCC-CM, MIROC5) under (A) Representative Concentration Pathway 4.5 (RCP 4.5), (B) RCP 8.5, and (C) combined under the two RCPs. Static land-use/land-cover projections were used for computation. Negative values represent a loss in mean habitat suitability between the two time periods, positive values a gain.

<u>Prediction unit:</u> Habitat Suitability Index (HSI) ranging from 0 to 1.

#### Uncertainty quantification

<u>Scenario uncertainty:</u> The uncertainty deriving from the use of four GCMs has been described in a spatially-explicit framework by computing the standard deviation among predictions from each GCM (Figure S1.7).

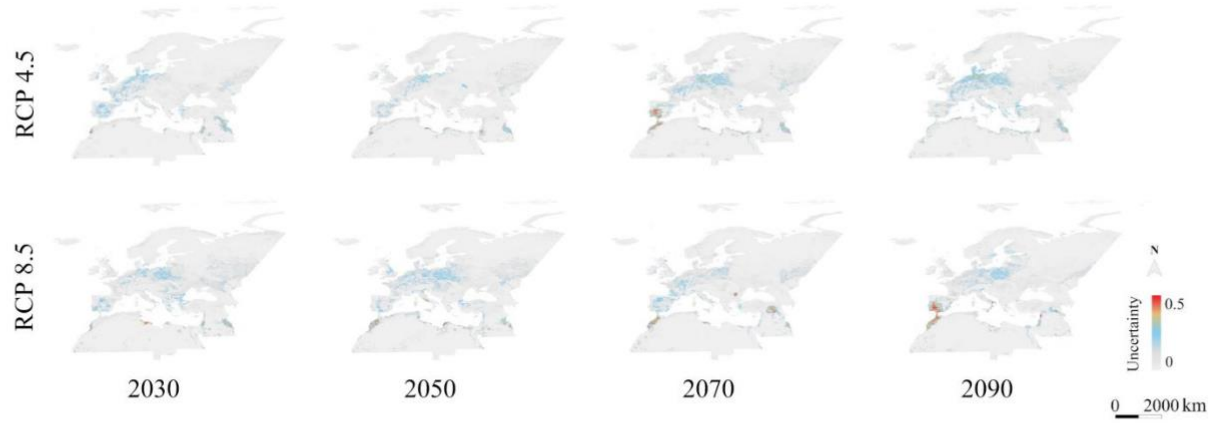


**Figure S1.7** Uncertainty resulting from the use of four General Circulation Models (GCMs) in static land-use/land-cover projections, expressed as the cell-wise standard deviation among habitat suitability projections under four GCMs (ACCESS1-3, CESM1-BGC, CMCC-CM and MIROC5).

<u>Novel environments:</u> We highlighted novel environments, i.e. areas where strict or combinational extrapolation occur, via the environmental overlap mask (Zurell *et al.,* 2012). Outputs from this analysis are found in Figures S1.8-S1.12.

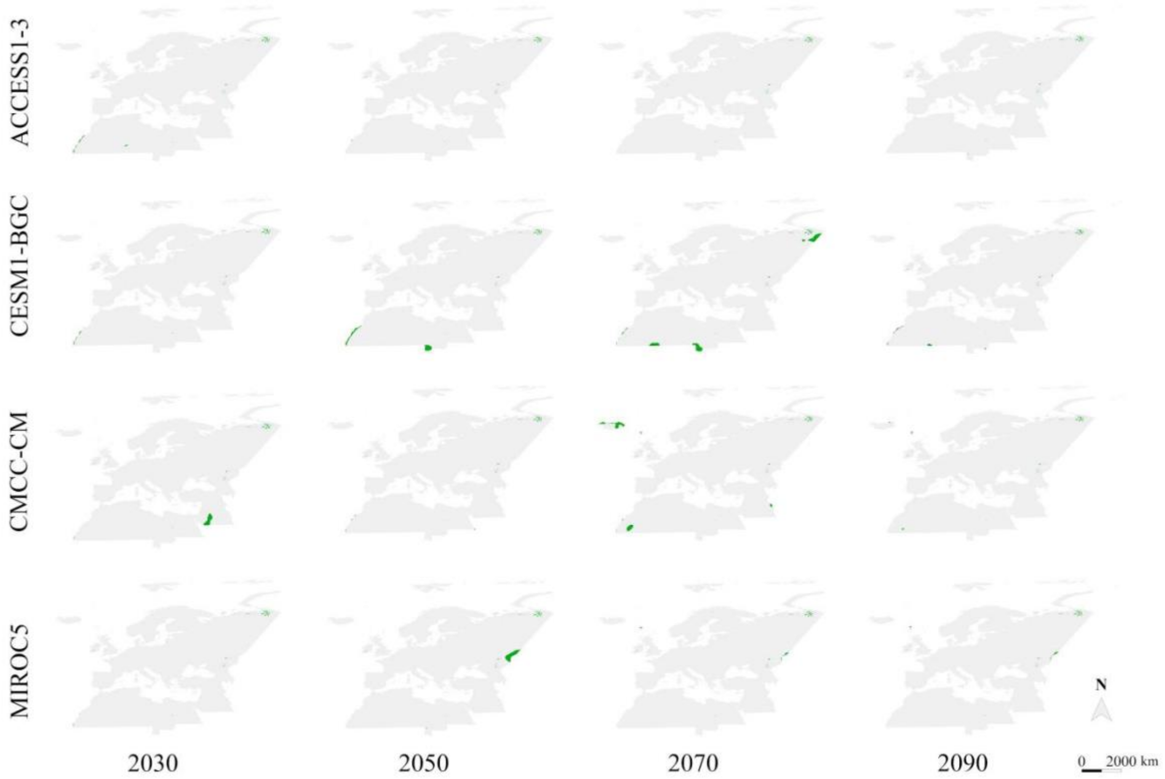


**Figure S1.8** Environmental overlap mask describing areas of strict extrapolation in static land-use/land-cover projections, under four General Circulation Models (ACCESS1-3, CESM1-BGC, CMCC-CM, MIROC5) and the Representative Concentration Pathway 4.5.

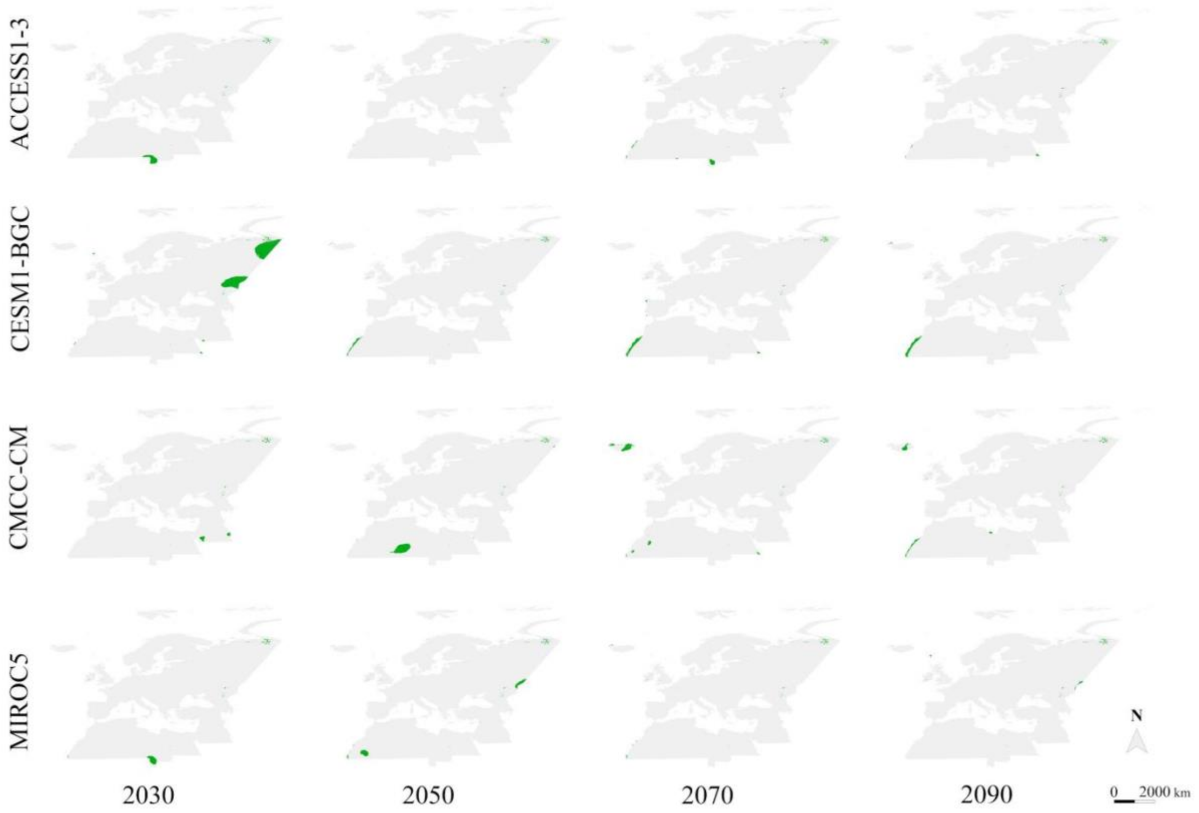


**Figure S1.9** Environmental overlap mask describing areas of strict extrapolation in static land-use/land-cover projections, under four General Circulation Models (ACCESS1-3, CESM1-BGC, CMCC-CM, MIROC5) and the Representative Concentration Pathway 8.5.

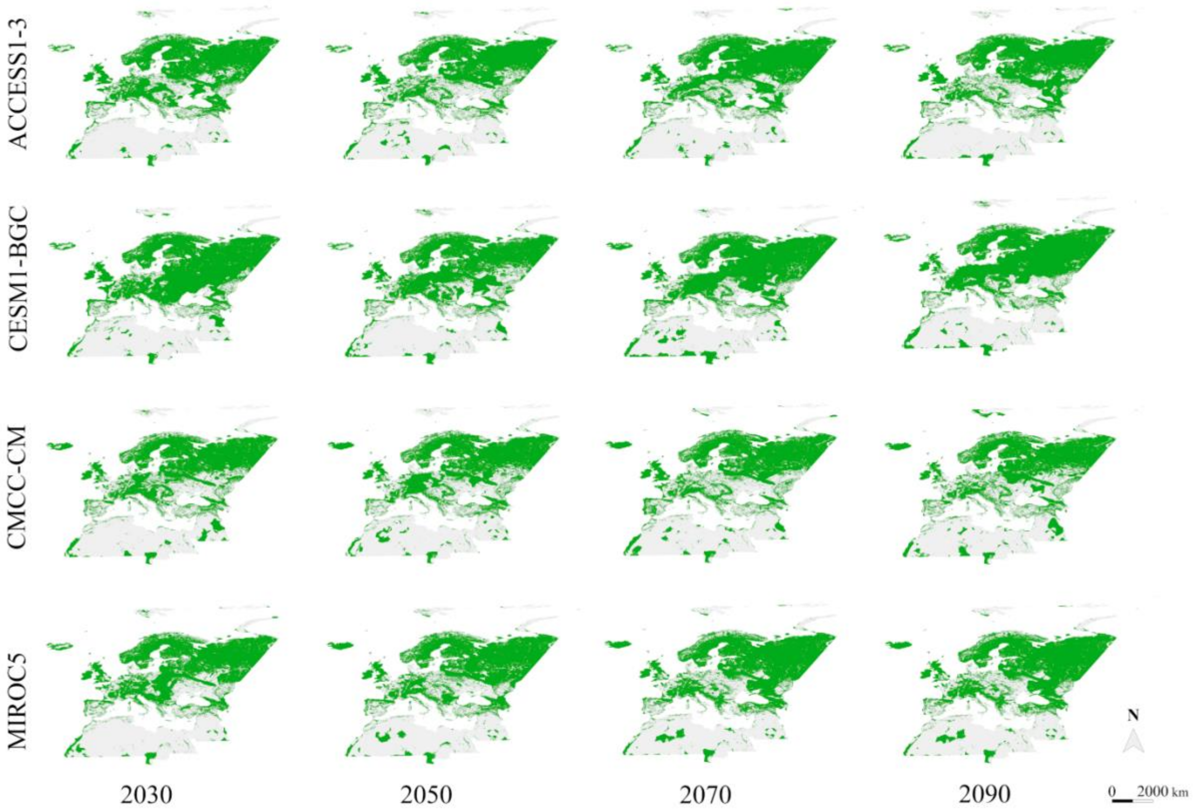


**Figure S1.10** Environmental overlap mask describing areas of both strict and combinational extrapolation in static land-use/land-cover projections, under four General Circulation Models (ACCESS1-3, CESM1-BGC, CMCC-CM, MIROC5) and the Representative Concentration Pathway 4.5.

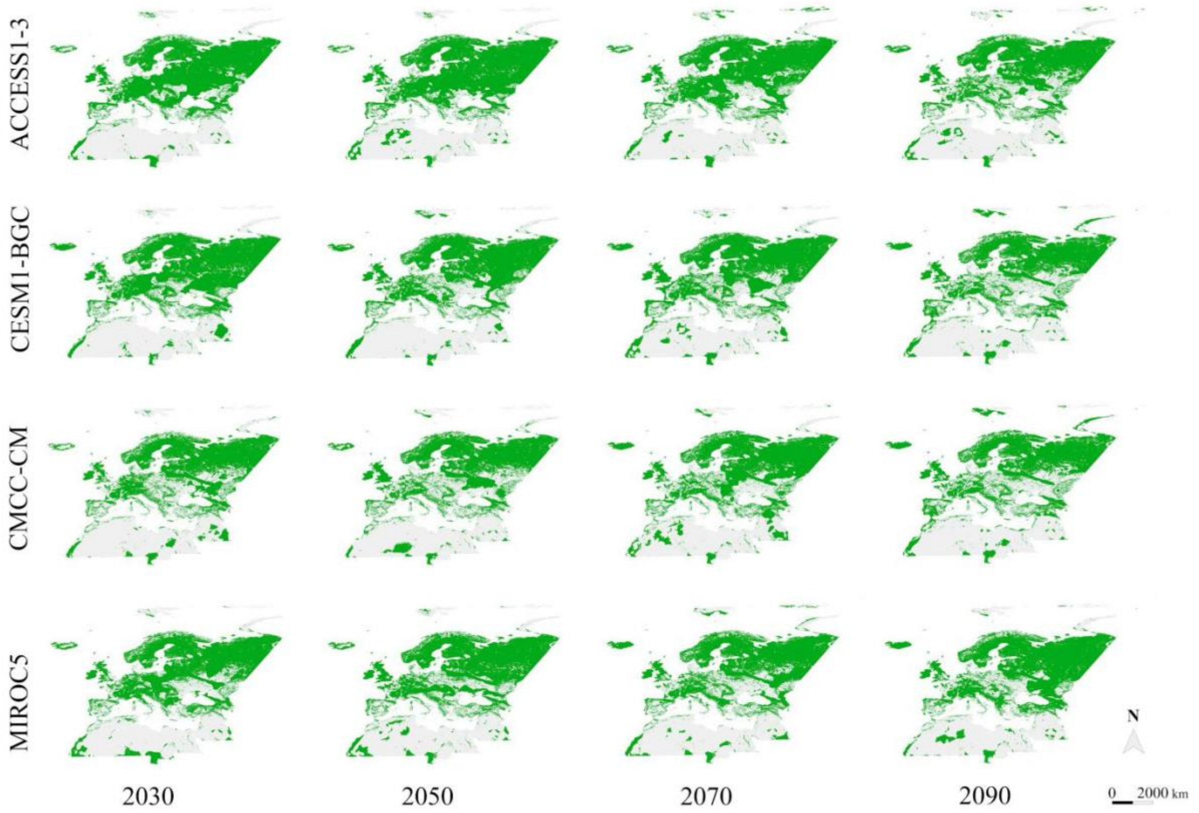


**Figure S1.11** Environmental overlap mask describing areas of both strict and combinational extrapolation in static land-use/land-cover projections, under four General Circulation Models (ACCESS1-3, CESM1-BGC, CMCC-CM, MIROC5) and the Representative Concentration Pathway 8.5.

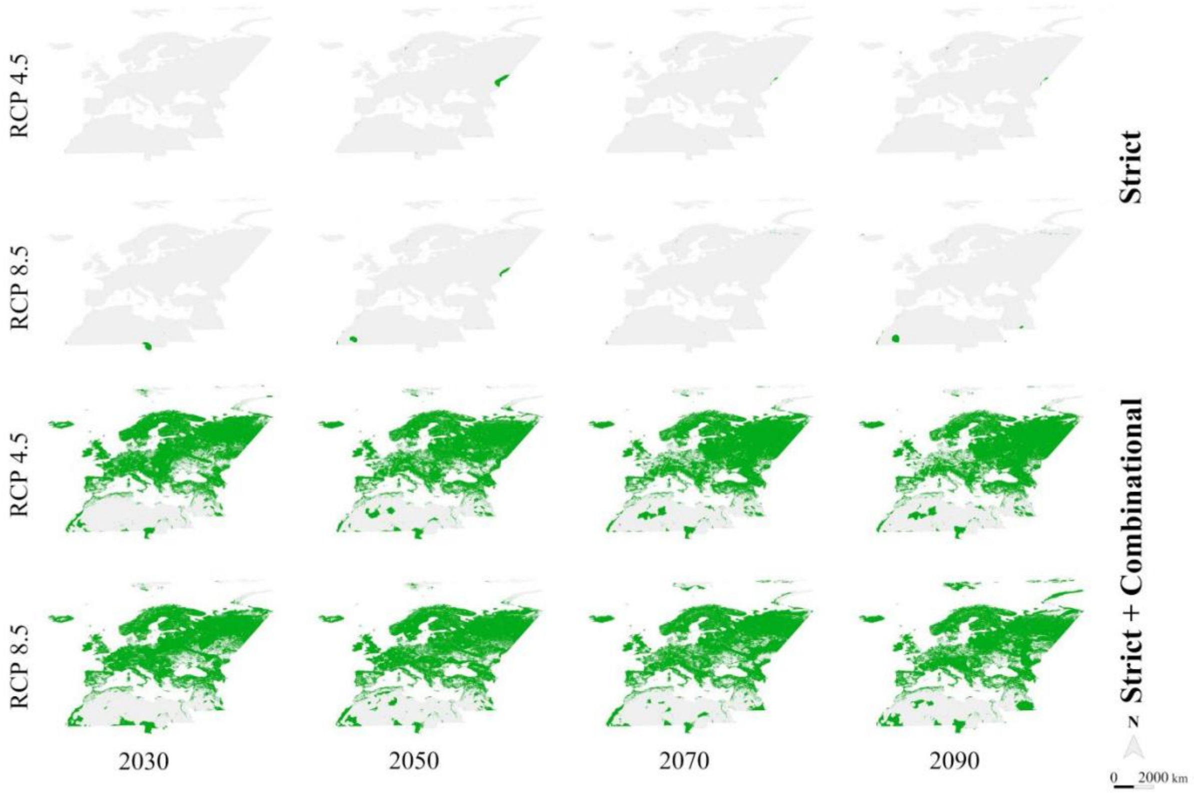


**Figure S1.12** Environmental overlap mask describing areas of strict or both strict and combinational extrapolation in dynamic land-use/land-cover projections, under the MIROC5 General Circulation Model, Shared Socioeconomic Pathway 5 and two Representative Concentration Pathways (RCP 4.5, RCP 8.5).
